# Endocytosis drives active cAMP signal discrimination among natively co-expressed GPCRs

**DOI:** 10.1101/2025.02.24.639927

**Authors:** Emily E Blythe, Rita R Fagan, Mark von Zastrow

## Abstract

Many G protein-coupled receptors (GPCRs) initiate a second phase of signaling after activation-induced endocytosis, but GPCRs vary considerably in their ability to internalize when activated. Here we show that this fundamental trafficking difference distinguishes the downstream signaling profiles of natively co-expressed GPCRs through the cAMP cascade. We focus on signaling to the nucleus stimulated by three different Gs-coupled GPCRs that are each endogenously co-expressed in human embryonic kidney cells but differ in their ability to internalize after activation: the adenosine-2B receptor that does not detectably internalize, the vasoactive intestinal peptide receptor-1 that internalizes very rapidly, and the β2-adrenergic receptor that internalizes less rapidly. We show that each GPCR produces a distinct signaling profile differentiated by endocytosis. Our results support a model in which endocytosis compresses chemical information sensed by distinct GPCRs into a spatiotemporal cAMP code by setting receptor-specific differences in the amount and duration of cAMP production from endosomes relative to the plasma membrane and that this is ‘decoded’ downstream in the pathway through sequential layers of processing by cytoplasmic and nuclear PKA activities. We propose that this biological information processing strategy has parallels to how computational encoder-decoder (autoencoder) networks denoise and recognize latent patterns in complex electrical signals.

## Introduction

Cells sense their chemical environment through membrane-embedded receptors that ultimately communicate the binding of a specific chemical ligand to the production of physiologically appropriate downstream functional responses. G protein-coupled receptors (GPCRs) comprise the largest and most diverse family of such chemical detectors in animal cells. GPCRs relay chemical information regulating downstream cellular physiology mainly through selective allosteric coupling among four different classes of heterotrimeric G proteins (*1*). Mammalian cells characteristically co-express a large repertoire of different GPCRs (e.g. (*2*)), enabling them to detect a chemical diversity far exceeding the number of available G protein classes. Nevertheless, there is remarkable physiological discrimination of ligand effects both in individual cells (*3–5*) and tissues (*6*). These facts make clear that downstream cellular ligand discrimination cannot be based solely on the inherent biochemical specificity of GPCR-G protein coupling, thus raising foundational questions about how different signaling profiles are generated in the presence of an effective information ‘bottleneck’ imposed on cellular biochemistry by the limited number of G protein classes available in the cell.

One strategy is to segregate GPCRs into different regions of the plasma membrane together with selected G protein-regulated effector proteins, effectively leveraging the plasma membrane as a two-dimensional surface for the lateral organization of specific signaling pathways (*1*, *7*). For example, in studies of signaling through the cAMP cascade, signaling specificity can be achieved through the lateral organization of nanometer-scale signaling domains stabilized by molecular scaffolds such as A-kinase anchoring proteins (AKAPs) (*8*, *9*) and through the inherent tendency of particular receptor and effector proteins to selectively self-assemble into homoligomeric complexes (*10*, *11*). A second strategy is to compartmentalize cellular signaling reactions in three dimensions using active trafficking processes to relocalize select signaling proteins to cellular endomembranes. Interest in this second strategy has intensified as considerable evidence has accumulated supporting the ability of GPCRs to undergo conformational activation by agonists in endomembranes as well as the plasma membrane and for the selective localization of various G proteins and G protein-regulated effectors to endomembranes that also contain active-conformation GPCRs (*12*).

Studies of cAMP signaling also provide strong support for the second strategy, demonstrating that various Gs-coupled GPCRs stimulate cAMP production in the cytoplasm and utilize endocytosis to promote downstream transcriptional control. A fascinating aspect of this transduction pathway is the long distance scale over which endocytosis-dependent effects must be communicated, as receptor-containing endosomes are typically orders of magnitude farther from the nucleus than the practical range of local cAMP concentration gradients (∼μm vs. ∼nm). It was initially proposed that endocytosis directs such distant downstream signaling simply by generating a sustained global cAMP elevation, based on studies of polypeptide hormone receptors that produce sustained responses to brief, pulsatile agonist application. It was later proposed that endocytosis selectively promotes a subset of effects, including signaling to the nucleus, after bath agonist application by changing the subcellular location of cAMP production, rather than its overall amount or duration (*12*). This concept of ‘location-encoded’ signaling from endomembranes, although initially proposed based on studies of catecholamine receptors that characteristically produce transient cAMP elevations (*13*, *14*), has since been supported from studies of signaling mediated by polypeptide hormone receptors (*15*, *16*). However, previous investigations have focused either on a single GPCR type that is endogenously expressed or on comparing signals produced from overexpressed receptors. It is not known if location-encoded signaling discriminates cellular signaling profiles generated through different GPCRs natively co-expressed after bath agonist application and, if so, it is not known how selectivity is communicated over a distance scale sufficient to impact signaling to the nucleus.

Here, we address both of these questions by focusing on signaling through the cAMP cascade stimulated by three distinct Gs-coupled GPCR types that are each endogenously co-expressed in HEK293 cells but differ in their ability to internalize after activation. Our results indicate that endocytosis does indeed distinguish the downstream signaling profiles among the natively co-expressed cellular GPCR complement and provide insight into a cellular information coding strategy sufficient to mediate long-range communication of selectivity. Based on these observations, together with accumulating mechanistic understanding of location-encoded signaling, we propose a model in which endocytosis drives an active biological signal processing scheme analogous to how computational encoder-decoder networks process and interpret complex electrical signals.

## Results

### β2AR, VIPR1, and A_2B_R are natively coexpressed but differ in their agonist-induced trafficking

As a first step toward testing the hypothesis that endocytosis enables cells to generate different native GPCR signaling profiles through the cAMP cascade, we sought to identify Gs-coupled receptors that are endogenously co-expressed in our HEK293 cell model and that are amenable to selective pharmacological activation with readily available drugs (Figure 1A). Previous studies have reported Gs-coupled signaling in HEK293 cells stimulated by the β-adrenergic catecholamine agonist Iso, the polypeptide hormone agonist VIP, and the adenosine analog 5’-N-ethylcarboxamidoadenosine (NECA) (*13*, *17*, *18*). We previously identified VIPR1 as the predominant GPCR mediating VIP-stimulated cAMP production our HEK293 system (*19*), and we employed pharmacological and genetic approaches to characterize the Iso- and NECA-sensitive GPCRs coexpressed in these cells. Using subtype-selective βAR antagonists (CGP 20712 for β1AR and ICI 118551 for β2AR) (*20*, *21*), we found the β2AR to be the predominant GPCR mediating Iso-stimulated cAMP cascade activation in the cells used in the present study (Supplementary Figure 1). NECA sensitivity in HEK293 has been previously attributed to expression of A_2B_R (*18*), and indeed, CRISPR knockout of A_2B_R largely abolished the NECA-stimulated cAMP response in our HEK293 system (Supplementary Figure 2A-B). Furthermore, we found that CGS-21680, a potent and selective agonist for the adenosine 2A receptor (A_2A_R), the other Gs-coupled adenosine receptor subtype, did not substantially stimulate cAMP production (Supplementary Figure 2C). Together, these results suggest that A_2B_R is the predominant GPCR mediating NECA-triggered activation of the endogenous cAMP cascade in our system.

**Figure 1.**
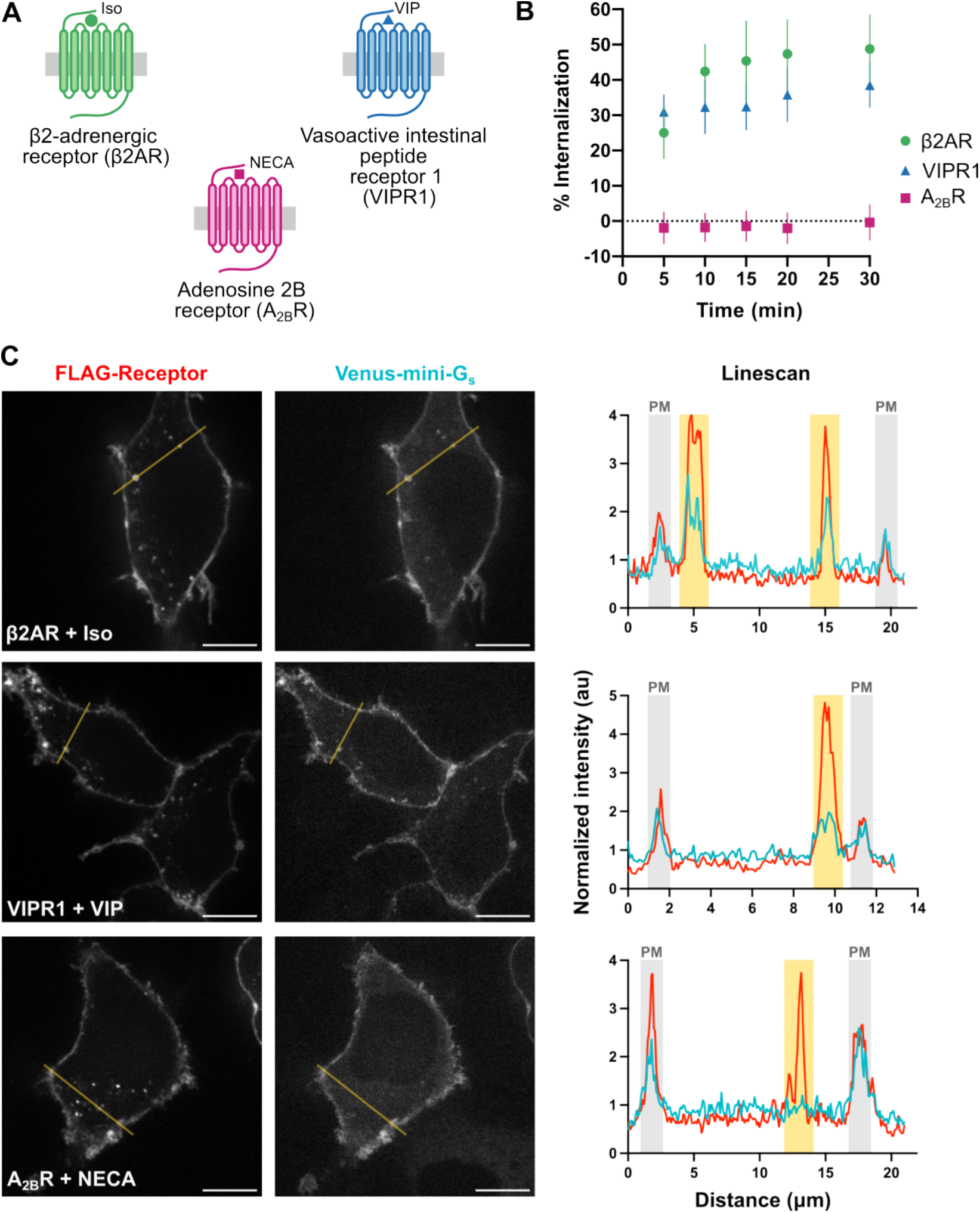
A_2B_R does not undergo appreciable agonist-induced endocytosis. (A) Three endogenously coexpressed GPCRs and their agonists used in this study. (B) Time course of agonist-induced internalization of HaloTagged GPCRs, as measured by flow cytometry. Receptors were stimulated with agonist (1 μM Iso, 500 nM VIP, or 20 μM NECA) for the noted times, and receptors remaining on the surface were labeled with a cell-impermeant HaloTag dye. N = 3 independent experiments, and error bars = S.D. (C) Colocalization of FLAG-tagged GPCRs and venus-mini-G_s_ 20 minutes after stimulation with agonist (1 μM Iso, 500 nM VIP, or 20 μM NECA). Prior to agonist stimulation, cells were treated with M1 anti-FLAG antibody labeled with Alexa Fluor 647. Scale bar is 10 μm. Linescans were normalized to the average intensity in each channel, with plasma membrane (grey) and receptor puncta (yellow) indicated. Representative images from N = 3 independent experiments.

We next asked if these GPCRs differ in their ability to undergo agonist-induced endocytosis. To do so, we used a flow cytometry-based internalization assay and a cell-impermeant HaloTag dye (JF_635_i-HTL) (*22*) to measure the effect of agonist application on plasma membrane levels of N-terminally HaloTagged receptors. Using this method, and consistent with previous reports (*17*, *19*, *23*), we observed very rapid agonist-induced internalization of the human VIPR1 and robust, but somewhat less rapid, internalization of β2AR (Figure 1B, note differences in time to steady state). We were unable to detect any internalization of the human A_2B_R over 30 minutes using the same Halo-tag assay system (Figure 1B), and similar results were obtained by an analogous method using a FLAG-tagged human A_2B_R (Supplementary Figure 3A). We were initially surprised by the lack of detectable A_2B_R internalization because a tagged rat A_2B_R was shown previously to rapidly internalize under similar conditions. However, this internalization process requires phosphorylation of a specific serine in the rat A_2B_R C-terminal tail (S329) (*24*) that is not conserved in the human A_2B_R (Supplementary Figure 3B), thus explaining the distinct trafficking behavior of the human A_2B_R.

Following internalization, β2AR and VIPR1 initiate a second phase of Gs-mediated signaling at the endosome (*19*, *25*). Since human A_2B_R does not detectably internalize, we anticipated that it produces less signaling from endomembranes than β2AR or VIPR1. To test this, we assessed receptor recruitment of mini-Gs (mG_s_), a G protein mimic that selectively binds active-conformation receptors (*26*, *27*). Venus-tagged mG_s_ was co-expressed with N-terminally FLAG-tagged receptors that were labeled at the cell surface using a fluorescent anti-FLAG antibody. For all three GPCRs, we observed agonist-dependent recruitment of mG_s_ to the cell periphery, indicative of agonist-induced conformational activation of receptors in the plasma membrane (Figure 1C). We also observed internal puncta of FLAG-labeled β2AR and VIPR1 in agonist-treated cells that colocalized with mG_s_, verifying the presence of conformationally activated receptors in endosomes as well as the plasma membrane (*19*, *25*). However, this was not the case for A_2B_R. Here, we observed FLAG-labeled intracellular puncta both in the presence and absence of NECA, suggesting some degree of constitutive (agonist-independent) internalization. However, none of the A_2B_R-labeled intracellular puncta colocalized with mini-G_s_. This suggests that the human A_2B_R signals from the plasma membrane but not from endosomes.

### Distinct profiles of endogenous β2AR, VIPR1, and A_2B_R signaling

Because the three endogenously coexpressed GPCRs differ both in agonist-induced internalization and the ability to accumulate in endomembranes in an active conformational state, we were next curious how these differences might affect downstream cellular signaling profiles signaling elicited by Iso, VIP, and NECA through the cAMP pathway. To characterize signaling at different stages with high temporal resolution, we employed genetically encoded real-time fluorescent sensors of cAMP (cADDis) and PKA activity (ExRai-AKAR2) (*28*). Importantly, these experiments were performed without overexpression of tagged receptors to measure signaling generated only by endogenous receptor pools. While we probed these two responses over a range of agonist concentrations (Supplementary Figure 4A), we focus in this section, for simplicity, on the effects of a saturating concentration of each corresponding agonist (Supplementary Figure 4B, Supplementary Table 1).

*Intracellular cAMP elevation*. All three agonists produced a similar peak level of global cAMP elevation detected by cADDis after bath agonist application (Supplementary Figure 5A), but each produced a different temporal profile (Figure 2A). The Iso response peaked and decreased gradually across the time course. The VIP response peaked and decayed very rapidly and then reached a second peak followed by a plateau, consistent with previous results (*19*). The NECA response differed still further, remaining largely prolonged at a high level throughout the measured time course of agonist exposure (Figure 2A). Confirming pronounced temporal differences in the cAMP responses produced by each agonist, we quantified decay constants and found them to vary over a range of ∼30-fold (Supplementary Table 2). We also calculated a sustained activity metric (*29*) that compares the signal remaining at 30 minutes post agonist addition to the peak signal. Using this metric, and fully consistent with the qualitative descriptions summarized above, we found that NECA elicited the most prolonged cAMP response, VIP elicited the most transient response, and Iso elicited a transient response but with overall slower kinetics than that elicited by VIP (Supplementary Figure 5B).

**Figure 2.**
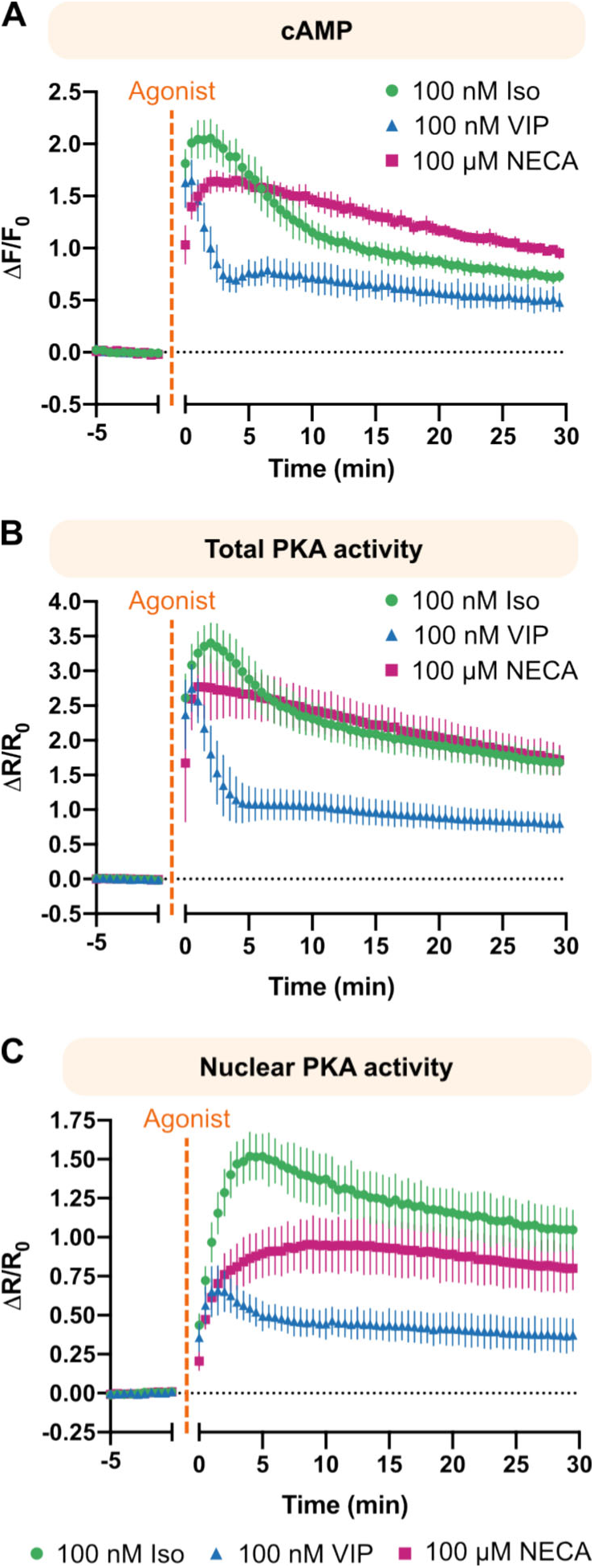
Endogenous GPCRs display unique cAMP and PKA signaling dynamics. Fluorescence time courses of biosensors for total intracellular cAMP (A), total cellular PKA activity (B), and nuclear PKA activity (C) upon stimulation of HEK293 cells with 100 nM Iso (green circles), 100 nM VIP (blue triangles), or 100 μM NECA (pink squares). N = 3 independent experiments, and error bars = S.D.

*Intracellular PKA activity.* We measured total intracellular PKA activity using ExRai-AKAR2, a well-established PKA activity biosensor (*28*). Each agonist produced a different temporal pattern of PKA activity elevation, but the overall dynamics of each response closely mirrored the respective global cAMP elevation measured using cADDis (Figure 2B). This close temporal correspondence was verified quantitatively by the similar metrics of peak and sustained determinations for each agonist between assays (Supplementary Figure 5A-B). Together, these results indicate that the global PKA activity elevations produced by each agonist are also different, but these differences largely follow differences in the dynamics of the global cellular cAMP elevation.

*Nuclear PKA activity.* To specifically probe the activity of PKA in the nucleus, we expressed a version of the PKA biosensor fused to a tandem nuclear localization signal (ExRai-AKAR2-2xNLS) (*30*) that enables transit through nuclear pores and drives efficient import from the cytoplasm. Each agonist produced a different profile of nuclear PKA activity elevation (Figure 2C). However, in contrast to the cytoplasmic PKA activity increase that generally mirrored the global cAMP elevation, the kinetics of the nuclear PKA activity increase differed markedly for all agonists. Specifically, the nuclear PKA activity rose more gradually than the cytoplasmic activity (Supplementary Figure 5C) and remained elevated for a longer period of time (Supplementary Figure 5B, Supplementary Table 2), consistent with previous studies (*31–34*) and the previously reported resemblance of nuclear PKA activity to a ‘low-pass’ filtered version of the cytoplasmic PKA activity that makes nuclear PKA activity selectively sensitive to the duration of upstream pathway activation (*31*).

### Effects of endocytosis on the endogenous GPCR signaling profiles

We next investigated how endocytosis impacts each of the measured signaling responses. As VIPR1 and β2AR internalize at different rates, and A_2B_R does not detectably internalize after activation, we anticipated that endocytic inhibition might PKA differentially affect the endogenous cellular responses elicited by each agonist. We inhibited endocytosis using a dominant negative mutant of dynamin (DynK44E), a manipulation known to block agonist-induced internalization of both VIPR1 and β2AR (*19*, *35*). We then assessed each signaling response in the same manner as above, normalizing each to its peak to focus specifically on the dynamics of signaling.

*Effect on global cAMP elevation.* Endocytic inhibition had little effect on the dynamics of the global cAMP elevation elicited by Iso or NECA, but it specifically abolished the second peak and plateau phase of cAMP elevation elicited by VIP (Figure 3A, Supplementary Figure 6). To estimate the relative contribution of endocytosis to the cAMP elevation, we calculated the difference between the control and DynK44E curves (Figure 3B). Consistent with previous results (*19*), this verified a substantial contribution of endocytosis to the global cAMP elevation elicited by VIP with a time course consistent with this component following VIPR1 internalization. In contrast, little endocytic contribution was evident in the Iso- and NECA-induced responses, consistent with β2AR stimulating significantly less cAMP production from endosomes than VIPR1 and A_2B_R barely internalizing. Using area under the curve (AUC) integration to estimate the overall endocytic contribution, we estimated a net endocytic contribution of ∼60% to the cAMP elevation elicited by VIP and ≤10% to that elicited by Iso or NECA (Figure 3C). Using the sustained activity metric, we verified a significant endocytic contribution to the VIP-elicited response but only a trend or no effect for Iso and NECA, respectively (Figure 3D). Thus, endocytic inhibition markedly suppressed the cAMP elevation elicited by VIP but had little effect on the Iso- or NECA-elicited responses, revealing that endocytosis indeed differentially sculpts the global cAMP responses produced by natively co-expressed GPCRs after bath agonist application.

**Figure 3.**
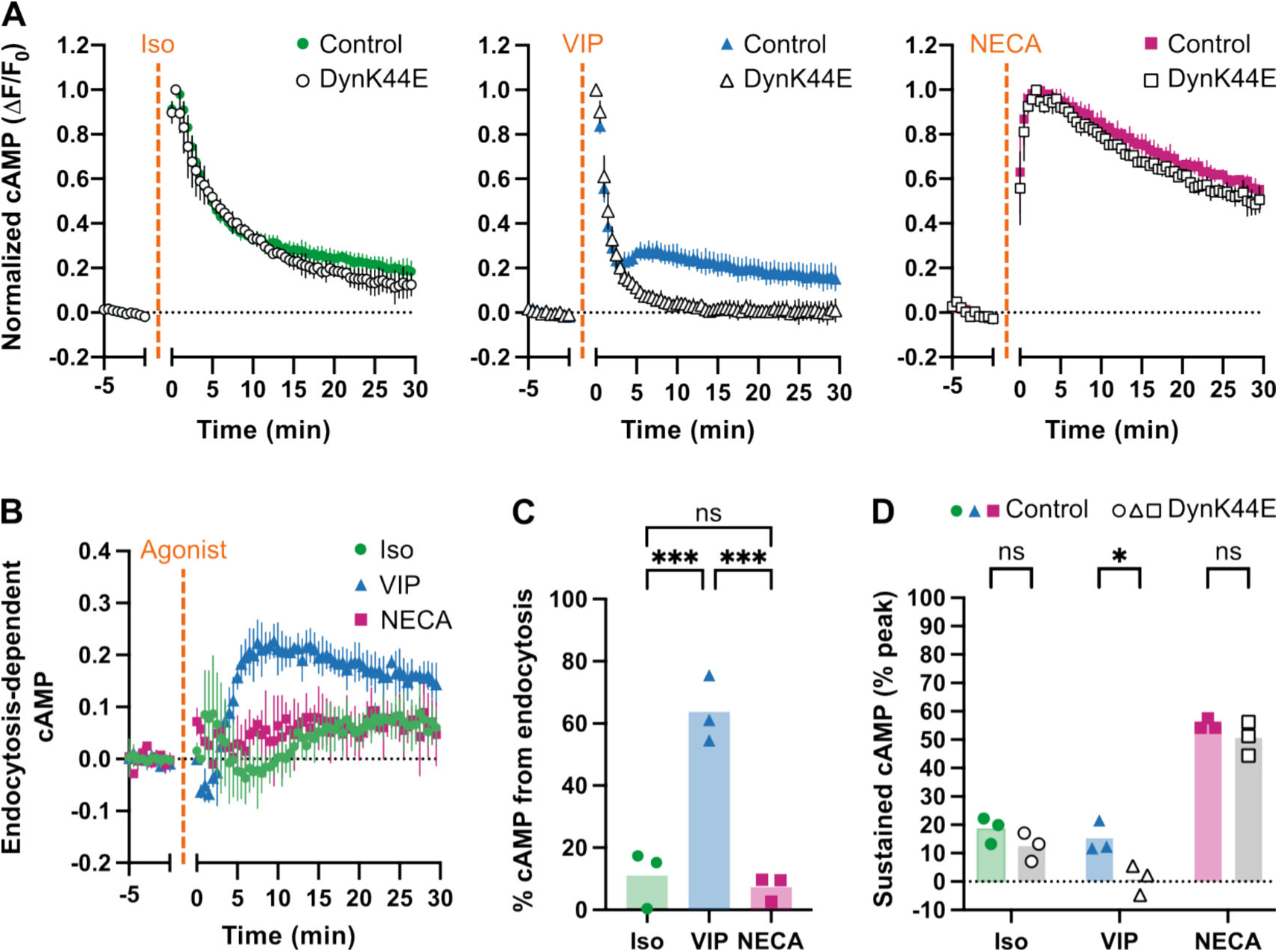
Endocytic inhibition halts VIP-stimulated cAMP signaling. (A) Fluorescence time course of a cAMP biosensor in cells expressing mCherry (control, colored shapes) or mCherry-DynK44E (DynK44E, open shapes) upon stimulation with 100 nM Iso (green circles), 100 nM VIP (blue triangles), or 100 μM NECA (pink squares). (B) Time course of the endocytosis-dependent fraction of cAMP responses, calculated as the difference between the control and DynK44E curves in (A). (C) Percent integrated cAMP signal that is dependent on endocytosis, calculated as the area under the difference curve in (B) normalized to the area under the corresponding control curve in (A). Significance was determined using an ordinary one-way ANOVA with Tukey’s multiple comparisons test. (D) Sustained cAMP signal 30 minutes after agonist addition, calculated as the percent fluorescence at 30 minutes from the curves in (A). Significance was determined using an ordinary two-way ANOVA with Sidak’s multiple comparisons test. For all panels, N = 3 independent experiments, and error bars = S.D.

*Effect on cytoplasmic PKA activity.* To specifically probe cytoplasmic PKA activity, we employed ExRai-AKAR fused to a nuclear export signal (ExRai-AKAR2-NES) (*30*). Endocytic inhibition markedly suppressed the cytoplasmic PKA activity increase elicited by Iso and VIP at later time points, but it had little or no effect on the NECA-elicited activity increase (Figure 4A, Supplementary Figure 7). This trend largely tracked the effects on global cAMP elevation for VIP (big effect) and NECA (no effect) but diverged strikingly for Iso (Figure 4B). Endocytosis enhanced the Iso-elicited increase of cytoplasmic PKA activity far beyond its effect on the global cAMP elevation, and this was verified by AUC integration (Figure 4C). The sustained activity metric (Figure 4D) further indicated that, for all agonists, the signal-promoting effect of endocytosis was on the later phase. Returning to the time course of the difference curves (Figure 4B), we also noted that the endocytosis-dependent component of cytoplasmic PKA activation elicited by Iso had a slower rate of onset than that elicited by VIP, consistent with β2AR internalizing less rapidly than VIPR1 (Figure 1B). Moreover, we noted that the endocytosis-dependent increase of the VIP-elicited PKA activity (average 7 min to peak) occurred with a faster rise time than the increase of global cAMP (average 9.8 min to peak; compare Figure 4B to Figure 3B). Together with previously established mechanistic information (*13*, *14*, *30*, *36*), these observations support the interpretation that endocytosis increases the efficiency of PKA activation elicited both by Iso and VIP, but not NECA, by moving the location of subcellular cAMP production into close proximity to concentrated PKA stores in the cytoplasm. They also provide additional support for the conclusion that endocytosis distinguishes the downstream signaling profiles of different GPCRs that are endogenously co-expressed in the same cells.

**Figure 4.**
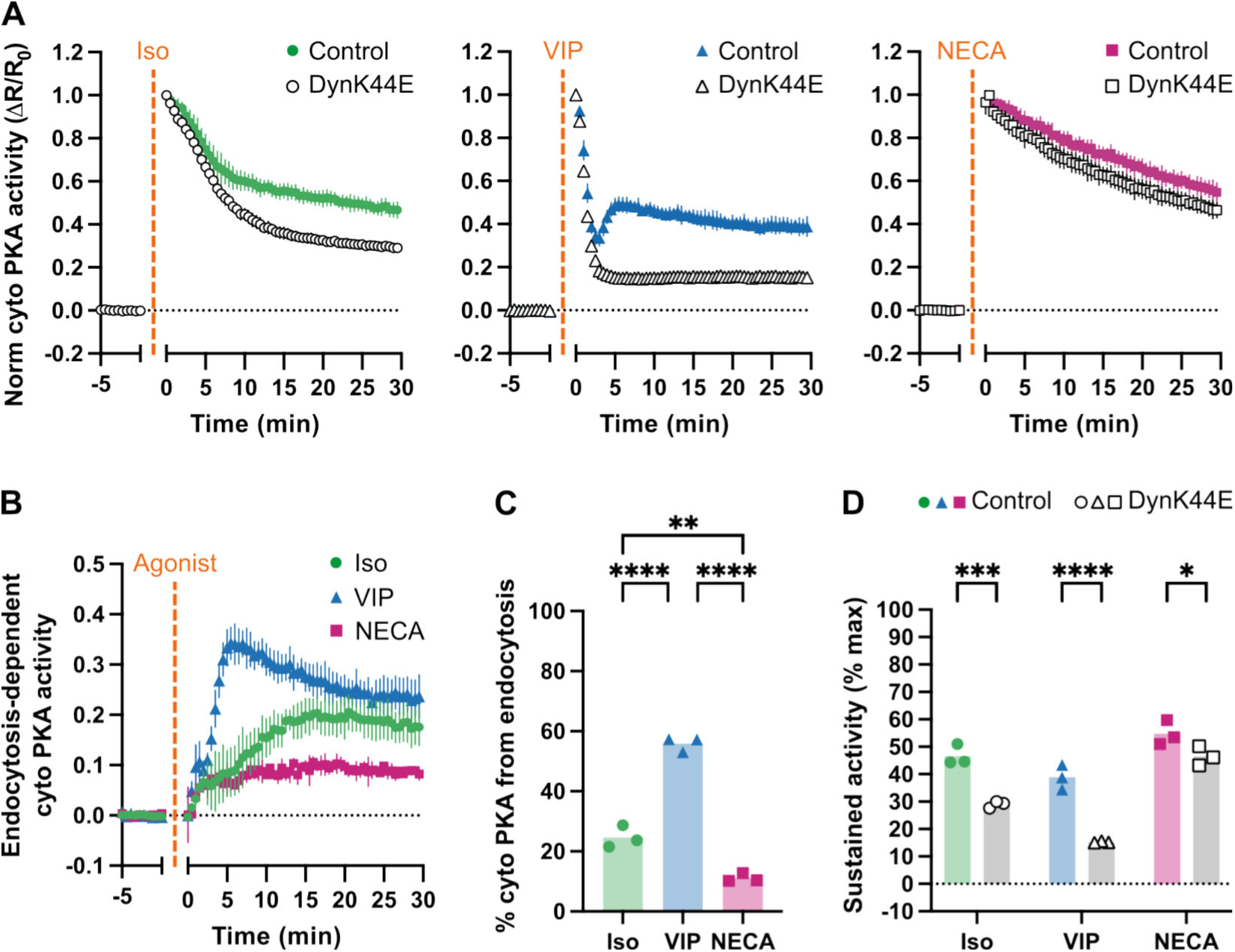
Endocytic inhibition dynamically blunts Iso- and VIP-stimulated cytoplasmic PKA activity. (A) Fluorescence time course of a cytoplasmic PKA activity biosensor in cells expressing mCherry (control, colored shapes) or mCherry-DynK44E (DynK44E, open shapes) upon stimulation with 100 nM Iso (green circles), 100 nM VIP (blue triangles), or 100 μM NECA (pink squares). (B) Time course of the endocytosis-dependent fraction of cytoplasmic PKA responses, calculated as the difference between the control and DynK44E curves in (A). (C) Percent integrated cytoplasmic PKA activity that is dependent on endocytosis, calculated as the area under the difference curve in (B) normalized to the area under the corresponding control curve in (A). Significance was determined using an ordinary one-way ANOVA with Tukey’s multiple comparisons test. (D) Sustained cytoplasmic PKA activity 30 minutes after agonist addition, calculated as the percent fluorescence at 30 minutes from the curves in (A). Significance was determined using an ordinary two-way ANOVA with Sidak’s multiple comparisons test. For all panels, N = 3 independent experiments, and error bars = S.D.

*Effect on nuclear PKA activity.* Significant differences were also observed in the effect of endocytic inhibition on the nuclear PKA activity increase (Figure 5A, Supplementary Figure 8). Here again, endocytic inhibition strongly suppressed the later phase of the response elicited by both Iso and VIP, but it had little or no effect on the response elicited by NECA (Figure 5B). AUC integration verified these agonist-selective effects (Figure 5C). The sustained activity metric further indicated a significant contribution of endocytosis to the late phase of nuclear PKA activity elevation elicited by Iso and VIP but essentially none to the elevation elicited by NECA (Figure 5D). However, while endocytosis clearly increased the magnitude of the nuclear PKA activity increase elicited by Iso and VIP (not NECA), the overall magnitude of this effect was similar to that measured at the level of cytoplasmic PKA activity (Figure 4C and 5C).

**Figure 5.**
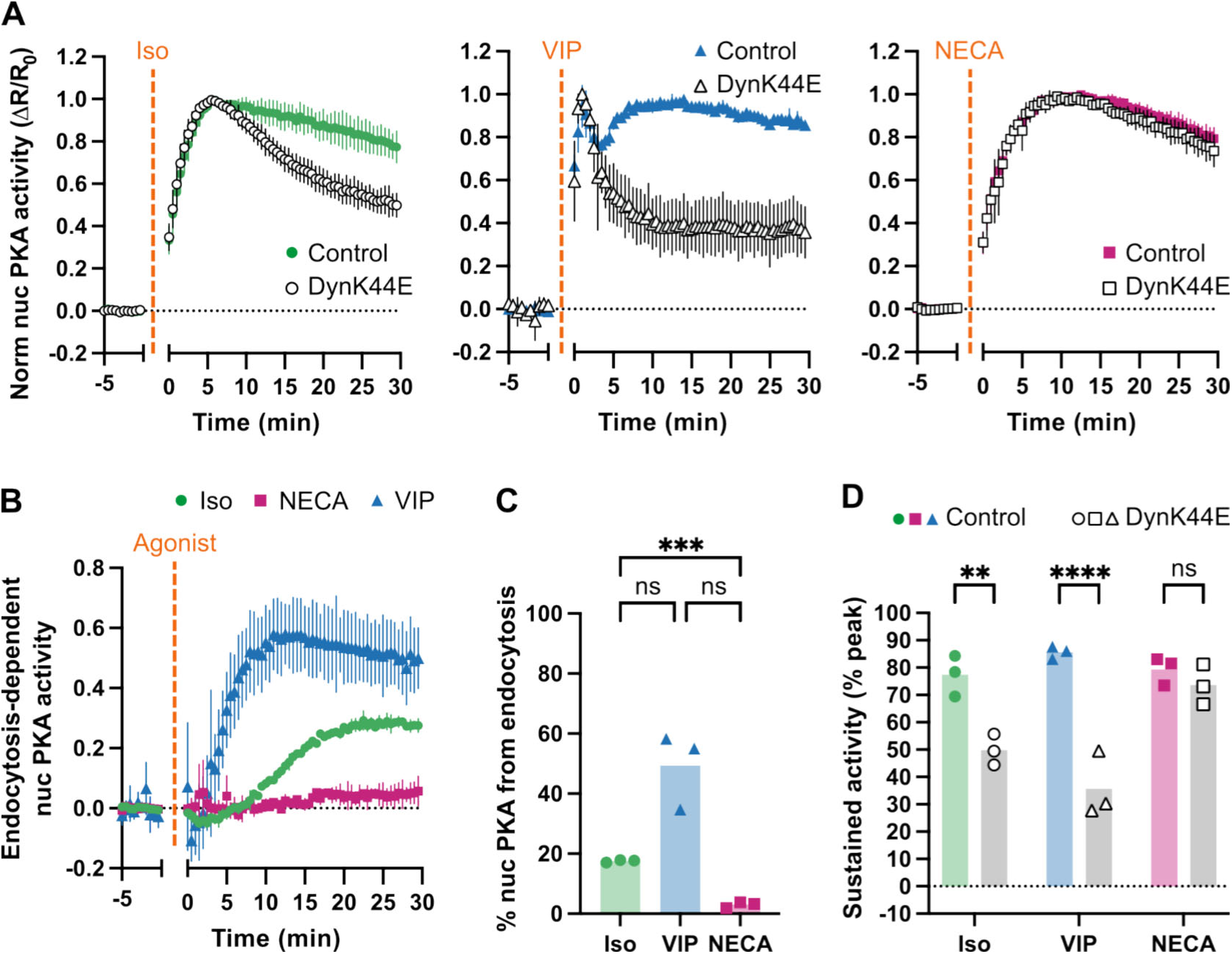
Only NECA can promote sustained nuclear PKA activity upon endocytic inhibition. (A) Fluorescence time course of a nuclear PKA activity biosensor in cells expressing mCherry (control, colored shapes) or mCherry-DynK44E (DynK44E, open shapes) upon stimulation with 100 nM Iso (green circles), 100 nM VIP (blue triangles), or 100 μM NECA (pink squares). (B) Time course of the endocytosis-dependent fraction of nuclear PKA response, calculated as the difference between the control and DynK44E curves in (A). (C) Percent integrated nuclear PKA activity that is dependent on endocytosis, calculated as the area under the difference curve in (B) normalized to the area under the corresponding control curve in (A). Significance was determined using a Brown-Forsythe ANOVA test with Dunnett’s T3 multiple comparisons test. (D) Sustained nuclear PKA activity 30 minutes after agonist addition, calculated as the percent fluorescence at 30 minutes from the curves in (A). Significance was determined using an ordinary two-way ANOVA with Sidak’s multiple comparisons test. For all panels, N = 3 independent experiments, and error bars = S.D.

Relationship to downstream control of cAMP-dependent transcription.

Finally, we examined cAMP-dependent transcriptional induction as a functionally relevant downstream signaling readout by measuring the stimulation of cAMP-dependent response element (CRE)-driven luciferase expression. We quantified the response by measuring luciferase activity after 5 hours of bath application of Iso, VIP or NECA, normalizing each measured response to that produced by direct activation of cellular adenylyl cyclases using forskolin. All of the agonists tested produced a robust, concentration-dependent transcriptional response by this metric (Figure 6A). Interestingly, NECA produced the strongest maximal response among the agonists tested (Figure 6B), despite its low potency (Figure 6A) and apparent failure to stimulate internalization of the human A_2B_R or detectable A_2B_R activation in endomembranes (Figure 1).

**Figure 6.**
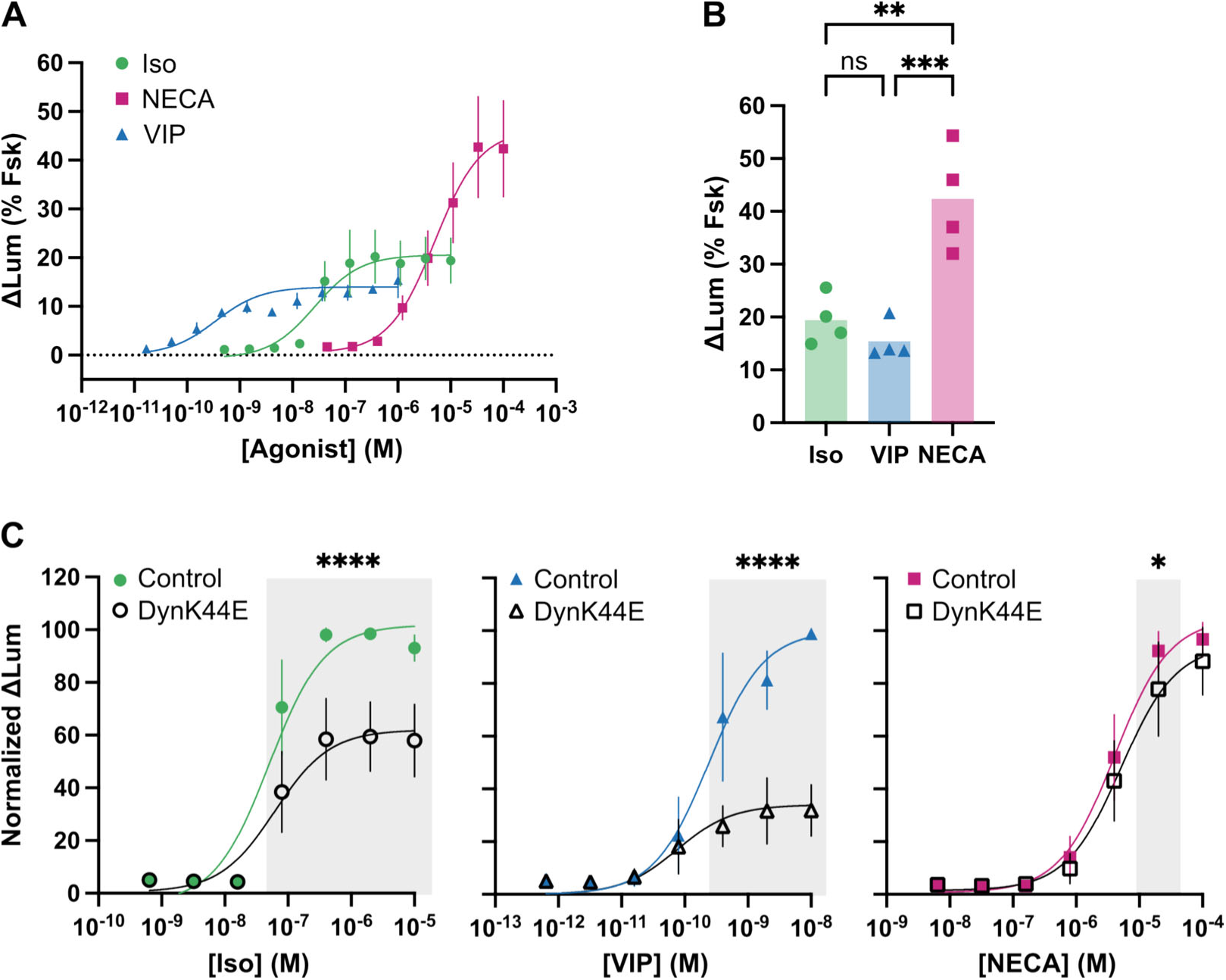
NECA-stimulated transcriptional activation is insensitive to endocytic inhibition. (A) Expression of CRE-luciferase transcriptional reporter upon treatment of cells with Iso (green circles), NECA (pink squares), or VIP (blue triangles) for 5 hours. Change in luminescence (ΔLum) was normalized to that of a forskolin-treated control within each replicate. Three-parameter dose-response curve fits are shown, and EC_50_s are listed in Supplementary Table 1. (B) Transcriptional reporter data from the highest agonist concentrations in (A). Significance was determined using an ordinary one-way ANOVA with Tukey’s multiple comparisons test. (C) Transcriptional reporter assays in cells expressing mCherry (control, colored shapes) or mCherry-DynK44E (DynK44E, open shapes). Cells were treated with Iso (green), VIP (blue), or NECA (pink) at the indicated concentrations for 5 hours. ΔLum was normalized to the maximum control ΔLum within each replicate. Three-parameter dose-response curve fits are shown, and EC_50_s are listed in Supplementary Table 1. Significance was determined using a two-way repeated measures ANOVA with Sidak’s multiple comparisons test. For all panels, N = 4 independent experiments, and error bars = S.D.

We then assessed the effect of endocytic inhibition imposed by DynK44E (Figure 6C). Endocytic inhibition suppressed the magnitude of the transcriptional response elicited by Iso and VIP by ∼40% and ∼60%, respectively, with little or no change in its concentration-dependence (Supplementary Table 1). In contrast, endocytic inhibition had very little effect on the transcriptional response elicited by NECA, reducing the measured luciferase activity by ≤15% and producing a statistically significant suppression at only one concentration. These observations verify that endocytosis has a strong downstream signal-promoting effect for β2AR and VIPR1 but not A_2B_R, tracking with differences in the ability of each GPCR to internalize after activation. They also reveal that A_2B_R robustly stimulates signaling to the nucleus despite the very weak ability of the human A_2B_R to undergo agonist-induced endocytosis. Accordingly, they indicate that differences among natively co-expressed GPCRs in their ability to internalize after activation, while significant, are not sufficient to fully explain the presently observed discrimination among the signal profiles of natively co-expressed GPCRs.

## Discussion

The present results answer the fundamental questions that we set out to address: First, does endocytosis confer selectivity on the downstream signaling profiles through the conserved cAMP cascade stimulated by GPCRs that are natively co-expressed and under conditions of simple bath agonist application? Second, if this is the case, how is it possible for endocytosis to communicate receptor-specific information to transcriptional control considering the long distance scale of signaling to the nucleus relative to local chemical gradients?

Our results answer the first question clearly in the affirmative, and they provide key insight into how distinct signaling profiles can be communicated over a long range in the cytoplasm. Overall, our results support a current understanding of location-encoded (or -biased) cellular GPCR signaling based on cAMP produced from endomembranes activating internal cytoplasmic PKA stores more efficiently than cAMP produced from the plasma membrane (*12*, *14*, *30*, *36*). They also show, we believe for the first time, that this principle is relevant to discriminating downstream cellular signaling profiles among distinct GPCRs that are natively co-expressed. Moreover, our results reveal additional complexity in cellular GPCR signaling not previously established or expected. First, we show that endocytosis enables native VIPR1 to produce a clear second phase of global cAMP elevation but, in the same cells, it enhances native β2AR signaling selectively downstream of global cAMP elevation. Second, while endocytosis has no detectable impact on the ability of A_2B_R to elevate cAMP or increase PKA activity, the native A_2B_R is nevertheless able to stimulate downstream transcriptional control robustly and in the same cell background in which both VIPR1 and β2AR produce clearly endocytosis-dependent transcriptional effects.

Our present interpretation of the above observations, taken altogether, is that the subcellular location of cAMP production is one important dimension differentiating the signaling profiles of natively co-expressed GPCRs, but it is not the only one. Rather, we propose that endocytosis functions in a cellular coding scheme that enables the long-range communication of GPCR-specific information based not only on the location, but also on the amount and duration of cytoplasmic cAMP production. In other words, endocytosis programs the spatial dimension of a spatiotemporal cAMP code that contains additional, ‘orthogonal’ information about amount and duration. The spatiotemporal coding hypothesis makes good sense in principle, as three dimensions are sufficient to describe the properties of any chemical signal, but it is uselessly vague in practice. In the following paragraphs, we propose a framework for achieving a more precise understanding of how distinct cellular signaling profiles are programmed, communicated, and interpreted through spatiotemporal cAMP coding.

Our proposal is based on a very simple concept of how GPCR signaling through cAMP is physically organized in cells by endocytosis (Figure 7A). We propose that the vast chemical diversity sensed through natively coexpressed GPCRs is ‘compressed’ into a spatiotemporal code in the cytoplasm by a process that is actively driven by endocytosis, first enabling GPCR-stimulated cAMP production from endomembranes in the first place and then sculpting differences in the amount and duration of production from endosomes relative to the plasma membrane in a receptor-specific manner. We further propose that this spatiotemporal code is subsequently decompressed or decoded through multiple layers of downstream processing that are configured effectively in series, with each layer providing different sensitivities to latent features of the code. A key layer sensitive to the location of cAMP production is cytoplasmic PKA activity, based on cAMP produced from endosomes being advantageously positioned for efficient activation of cytoplasmic PKA due to local proximity to concentrated PKA stores (*14*). We propose that nuclear PKA activity functions downstream of this layer and provides additional decoding. First, it effectively ‘passes through’ spatial information decoded upstream by cytoplasmic PKA, and then it confers additional sensitivity to the duration of pathway activation that is largely irrespective of subcellular location (*31*). This process of multi-layered decoding is inherently complex because it is quantitative rather than qualitative and involves the passage of information through different layers, each of which is sensitive to more than one dimension of the cAMP code but to different relative degrees.

**Figure 7.**
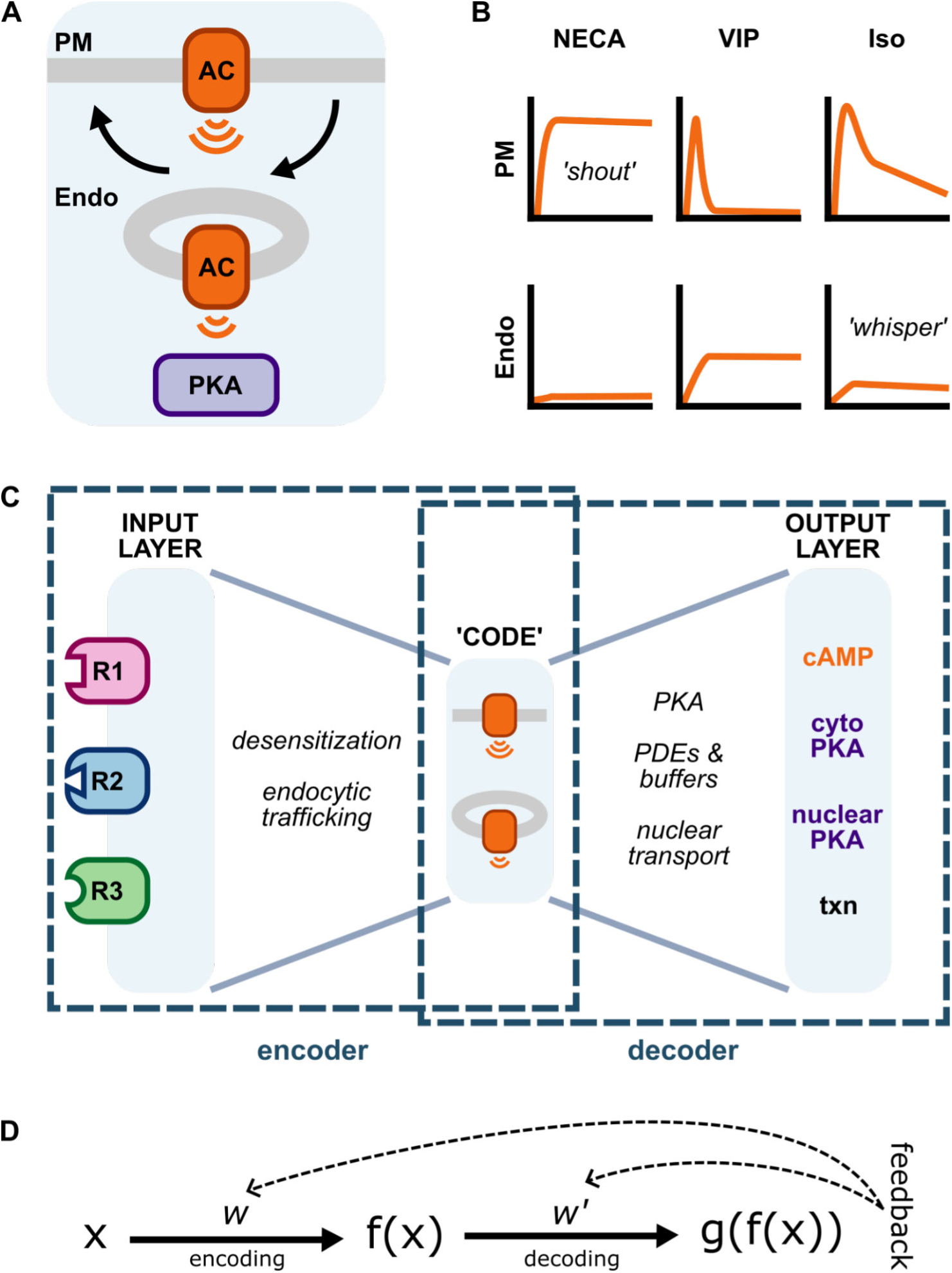
Signaling through a spatiotemporal cAMP code. (A) Adenylyl cyclases (ACs) generate cAMP from both the plasma membrane (PM) and endosomes (endo). GPCRs produce distinct codes through differences in the amount and duration of cAMP production they stimulate from each location, defined by receptor-specific distinctions in desensitization and trafficking properties. (B) NECA triggers native A_2B_R to stimulate prolonged cAMP production from the PM (top curve) but almost none from endosomes (bottom curve). VIP triggers native VIPR1 to stimulate transient cAMP production from the PM and prolonged production from endosomes. Iso triggers native β2AR to stimulate transient cAMP production from the PM more strongly than VIP, and prolonged production from endosomes less strongly than VIP. The cAMP code contains latent features that are then interpreted through multiple ‘layers’ of post-processing, as discussed in the text. (C) The integrated ‘encoder-decoder’ scheme that we propose, emphasizing its similarity to a computational autoencoder. (D) A more simplified version, showing a proposed error correction strategy by feedback gradient back-propagation. X represents the input vector, f(X) represents its compressed version in the cAMP code layer, and g(f(X)) represents the decompressed or decoded version used to control downstream signaling effects. Adjustable parameter matrices fine-tuning encoding and decoding operations are represented by *w* and *w’*, respectively.

The information processing strategy proposed above is certainly an oversimplification, and it does not include a number of mechanistic elements that are well-known to influence cellular cAMP signaling. We suggest that these may be considered additional layers of processing beyond those discussed here. Nevertheless, our model comports with long-recognized core features of cAMP signaling through PKA (*37*, *38*) and with more recent evidence indicating that cAMP production from endosomes promotes the entry of free PKA catalytic subunits into the nucleus from the cytoplasm (*14*). Thus we suggest that the present model, while admittedly simplistic, provides a reasonable starting point for developing an integrated understanding.

Experimental observations supporting our proposed model are summarized schematically in Figure 7B. NECA triggers A_2B_R to stimulate prolonged cAMP production from the PM (top panels) but almost no production from endosomes (bottom panels). VIP triggers VIPR1 to stimulate transient cAMP production from the PM and prolonged production from endosomes. Iso triggers β2AR to stimulate transient cAMP production from the PM more strongly than VIPR1 and prolonged production from endosomes less strongly than VIPR1. These differences program quantitative differences that accordingly define a quantitative spatiotemporal code of receptor-specified cAMP production in the cytoplasm whose dimensions determine, collectively, downstream functional effects. NECA elicits a prolonged, high level of cAMP production from the plasma membrane that robustly drives all downstream effects, including transcriptional control. As a simple metaphor, prolonged cAMP production from the plasma membrane resembles a ‘shout’ that broadly instructs downstream effects independently of endocytosis. Iso elevates global cAMP transiently and elicits only a small amount of cAMP production from endosomes. This is not sufficient to detectably increment the global cAMP elevation, but the cAMP production from endosomes is prolonged due to iterative receptor cycling in the presence of agonist and activates cytoplasmic PKA efficiently by proximity. Thus the endosome can effectively ‘whisper’ instructions to drive transcriptional effects without changing global cAMP. The receptor-specific code is then decompressed or decoded quantitatively in layers, using cytoplasmic PKA activity to selectively sense location and nuclear PKA activity in a more downstream processing layer to pass through spatial information detected upstream and additionally sense the temporal dimension, based on the previously established low-pass filtering properties of nuclear PKA activity that make it inherently sensitive to the duration of activation (*31*).

Viewed more broadly, we speculate that this model of spatiotemporal cAMP encoding-decoding has parallels to the operation of an ’autoencoder’ network of the sort widely deployed in machine learning and electronic signal processing (*39*) (Figure 7C). We suggest that this analogy is useful because computational autoencoders, despite relying on quantitative processing of information after compression into an abstract internal code, have been proven to be highly efficient tools for denoising and recognizing latent features in complex electrical signals. An autoencoder is composed of two key parts: an ‘encoder’ that compresses a complex input into an internal code layer of reduced dimensionality, and a ’decoder’ that decompresses this code into a filtered, denoised version of the input or a transformed version highlighting latent features. We speculate that cellular GPCR signaling mediated by the cAMP cascade operates in a similar way. Here, the complex chemical environment sensed by the co-expressed GPCR cohort (R1-3 in the diagram) defines the input, and this is compressed into a spatiotemporal code defined by the amount, duration and location of cAMP production in the cytoplasm. Encoding is mediated primarily through differences in receptor desensitization and trafficking properties, resulting in each chemical input programming a quantitatively different spatiotemporal code. Decoding involves multiple layers of data transformation (often called ‘hidden layers’ in machine learning) that are effectively configured in series, each differing in quantitative sensitivity to distinct features of the code. As noted above, our results suggest that cytoplasmic PKA senses location and nuclear PKA passes through this information and additionally senses overall duration of upstream cascade activation. Like processing by an electronic autoencoder, however, no individual layer uniquely senses any single feature and all layers function together.

Extending the autoencoder analogy, a key outstanding question concerns how error correction is implemented in the network. The efficient operation of a computational autoencoder critically depends on proper adjustment of parameters that control the respective encoding and decoding operations (*40*) (Figure 7D). Typically, these ‘weights’ are first adjusted by a human programmer to set them in a reasonable range and then the network is fine-tuned by extensive ‘training’ sessions using reference data, in which an error gradient between the computed and desired network output is calculated across successive trials and ‘back-propagated’ through the network mathematically. For cellular signaling through the cAMP cascade, we speculate that the analogous weights are first set in a reasonable range by evolution, based on cell type-specific expression and localization of network components. We further speculate that fine-tuning is guided by feedback from target cells in tissues, enabling a form of self-correction instructed by downstream physiology, and that error gradients are calculated and back-propagated biochemically. This is plausible because cellular signaling reactions inherently generate spatial and temporal gradients, and it is already known that signaling can be modified by downstream PKA activity all the way back to regulatory phosphorylation of the GPCRs that constitute the input layer. Examples include the process traditionally called heterologous desensitization, in which PKA-mediated phosphorylation exerts negative feedback on GPCR signaling activity (*41*), and evidence that phosphorylation may switch its ‘sign’ from stimulatory to inhibitory in some cases (*42*, *43*).

In closing, the present study establishes an essential function of endocytosis in distinguishing the signaling profiles of natively co-expressed GPCRs through the cAMP cascade in intact cells, at fully endogenous levels of expression and after simple bath agonist application. To our knowledge, this is the first study to explicitly demonstrate such selectivity with all of these conditions met. In addition, our observations support an extensible framework on which to build an improved, systems-level understanding of signaling selectivity.

## Materials and Methods

### Cell Culture

HEK293 cells (ATCC CRL1573) were cultured in DMEM (Gibco 1196511) supplemented with 10% FBS (Hyclone SH30910.03 or R&D Systems S12495 via UCSF Tissue Culture Facility) and grown at 37 °C with 5% CO_2_. Halo-β2AR and inducible Halo-VIPR1 stable cell lines have been previously described (*19*). Cells stably expressing Halo-A_2B_R were selected with 25 μg/mL zeocin, and cells stably expressing FLAG-β2AR or FLAG-A_2B_R were selected with 500 μg/mL geneticin.

### CRISPR knockout of A_2B_R

Cells were electroporated with single guide RNAs (A_2B_R: GACACAGGACGCGCTGTACG, NT: GCACUACCAGAGCUAA; Synthego) in complex with Cas9 (UC Berkeley Macrolab) before monoclonal selection. Genetic modifications were verified by Sanger sequencing of PCR amplicons (primers: TGCGTGAGCACCAGCACGAA and GGCAATTTGTTAGTTATCCGCCGC).

### Reagents and antibodies

Information about the chemicals and antibodies used in this study can be found in Supplementary Tables 3 and 4.

### DNA Constructs and BacMam virus

Details about the plasmids and BacMam viruses used in this study can be found in Supplementary Table 5. Transfections using Lipofectamine 2000 (ThermoFisher) were carried out 24-48 hours before experiments according to the manufacturer’s protocol. BacMam viruses from pCMV-Dest plasmids were produced according to the manufacturer’s protocol (Invitrogen) and added to cells 24 hours before experiments.

### Fluorescent biosensor assays

Cells treated with cADDis (Montana Molecular #U0200G), ExRai-AKAR2, ExRai-AKAR2-NES, ExRai-AKAR2-2xNLS, mCherry, and/or mCherry-DynK44E BacMams were plated into 96-well plates (Corning 3340) coated with poly-L-lysine (Millipore Sigma P8920). After 24 h, cells were washed twice with assay buffer (20 mM HEPES pH 7.4, 135 mM NaCl, 5 mM KCl, 0.4 mM MgCl_2_, 1.8 mM CaCl_2_, 5 mM d-glucose) before a 10-min incubation in a prewarmed 37 °C plate reader (Spark, Tecan Life Sciences, controlled by SparkControl v3.2). When appropriate, 30 μM Dyngo4a or vehicle was added immediately prior to preincubation. For βAR antagonist experiments, cells were preincubated for 30 minutes in assay buffer containing antagonist (300 nM GCP 20712, 300 nM ICI 118551, or 10 μM Alp) or vehicle.

For cADDis, fluorescence was read at an excitation wavelength of 500/5 nm and an emission wavelength of 530/10 nm. For ExRai-AKAR2 constructs, fluorescence was read using 400/20 nm and 485/20 nm excitation filters and a 520/10 nm emission filter. mCherry was read using a 560/20 nm excitation filter and a 620/10 nm emission filter. Data were collected every 30 seconds for 35 minutes, with vehicle or agonist added at 5 minutes.

Intracellular cAMP was calculated using raw fluorescence (F), while intracellular PKA activity was calculated using the ratio of fluorescence (R = F_485_ / F_400_) background corrected by subtracting the average fluorescence of untransduced wells at each timepoint (*29*). Changes in cAMP (ΔF/F_0_) and PKA activity (ΔR/R_0_) were calculated as the change in fluorescence (ΔF = F − F_0_ or ΔR = R − R_0_) normalized to the average of the 5-min baseline (F_0_ or R_0_). For samples expressing mCherry or mCherry-DynK44E, curves were further normalized to the peak fluorescence change (ΔF/F_0_)_max_ or (ΔR/R_0_)_max_ as noted in figure legends. Difference curves were calculated by subtracting mCherry-DynK44E curves from mCherry curves. Sustained activity, measured 30 minutes post-agonist addition, was calculated as (ΔF/F_0_)_30_ / (ΔF/F_0_)_max_ x 100 or (ΔR/R_0_)_30_ / (ΔR/R_0_)_max_ x 100. Integrated signal was calculated as an area under the curve following agonist addition. For dose–response curves, data were fit to a three-parameter dose-response equation in Prism (v.10). When top and bottom plateaus could not be automatically fit, these were constrained to the average maximum signal and zero, respectively.

### Internalization by flow cytometry

Cells stably expressing N-terminally tagged GPCRs in 12-well plates were treated with Iso (100 nM or 1 μM), 500 nM VIP, or 20 μM NECA for the indicated times. After three washes with cold PBS, cell surface receptors were labeled with 200 nM JF_635_i-HTL (HaloTagged receptors) or M1 antibody conjugated to Alexa Fluor 647 (FLAG-tagged receptors) for 30 minutes at 4 °C. FLAG-tagged samples were mechanically lifted, while HaloTagged samples were washed once with PBS-EDTA (UCSF Cell Culture Facility) before being lifted with TrypLE Express (Thermo Fisher). Surface staining was measured on an Attune NxT Flow Cytometer equipped with a CytKick Autosampler and controlled by Attune Cytometric Software (v5.3.2415.0) using a 561 nm excitation laser and a 620/15 nm emission filter. Data were analyzed using FlowJo v.10.10.0 software (BD Life Sciences). Populations were gated for cells expressing receptor, and internalization was calculated as [1 - (F_t_ / F_t0_)] * 100, where F_t_ is the median fluorescence at time t. Each biological replicate represents the average of three technical replicates.

### Luminescence transcriptional reporter assay

Cells were transfected with pGL4.29[luc2P/CRE/Hygro] (Promega E8471) and replated into white 96-well plates (Corning 3917) coated with poly-L-lysine (Millipore Sigma P8920) at a concentration of 50,000 cells/well. If cells were pretreated with mCherry or mCherry-DynK44E, BacMam virus was added upon replating. After overnight incubation, media was replaced before treatment with vehicle (water or DMSO), Iso, VIP, or NECA at the indicated concentrations. Cells were incubated at 37 °C with 5% CO_2_ for 4.75 hours followed by a 15 minute incubation at room temperature. After a wash with room temperature assay buffer (20 mM HEPES pH 7.4, 135 mM NaCl, 5 mM KCl, 0.4 mM MgCl_2_, 1.8 mM CaCl_2_, 5 mM d-glucose), cells were treated with 1.6 mM D-luciferin (Gold Biotechnology LUCNA) in assay buffer. Luminescence was read two minutes later by plate reader (Spark, Tecan Life Sciences, controlled by SparkControl v3.2). Change in luminescence (ΔLum) was calculated as the ratio of the average luminescence of agonist-treated wells divided by the average luminescence of vehicle-treated wells. Within each biological replicate, changes in luminescence were normalized to either a 10 μM forskolin control or each agonist’s maximum response in the mCherry control.

### Live-cell imaging

Cells were cotransfected with FLAG-tagged GPCRs and venus-mini-G_s_ and replated into poly-L-lysine (Millipore Sigma P8920) coated glass bottom dishes (Cellvis D35-20-1.5-N). After 48 hours, cells were labeled with M1 antibody conjugated to Alexa Fluor 647 for 10 minutes at 37 °C, washed three times, and imaged live in DMEM without phenol red (Gibco 31053028) supplemented with 30 mM HEPES pH 7.4. Confocal microscopy was carried out on Nikon Ti inverted microscope controlled by NIS Elements HC v.5.20.02 (Nikon) and fitted with a CSU-22 spinning disk unit (Yokogawa), custom laser launch (100 mW at 405, 488, 561 and 640 nm, Coherent OBIS), Sutter emission filter wheel, an Apo TIRF ×100/1.49 numerical aperture oil objective (Nikon), and a Photometrics Evolve Delta EMCCD camera. Cells were kept in a temperature- and humidity-controlled chamber (Okolab) at 37 °C. Cells were imaged for 2 minutes before addition of agonist and 20 minutes after agonist. Images were processed and analyzed using Fiji (v.2.3.0) (*44*).

### Data analysis and reproducibility

Statistical analysis and curve fitting was carried out using Prism (v.10, GraphPad) as noted in figure legends. All data are shown as individual biological replicates or as mean ± standard deviation from at least three biologically independent experiments, unless otherwise noted. For fluorescent biosensor assays, luminescent transcriptional reporter assays, and flow cytometry internalization assays, each biological replicate represents the average of at least two technical replicates.

## Acknowledgements

We thank Jin Zhang and Luke Lavis for generously sharing reagents, Nicole Fisher for assistance with cloning and viral production, Uygar Sümbül and members of the von Zastrow lab for helpful discussion, and the following core facilities for providing services to support this research: the UCSF Center for Advanced Light microscopy (Nico Sturrman, Kari Herrington, Micaela Lasser, DeLaine Larsen, and SoYeon Kim) and the UCSF Helen Diller Family Comprehensive Cancer Center Laboratory for Cell Analysis (Sarah Elmes; supported by NIH under award P30CA082103).

## Author Contributions

E.E.B. and M.v.Z. conceived of the study. E.E.B., R.R.F., and M.v.Z. designed experiments. E.E.B. and R.R.F. carried out experiments. E.E.B., R.R.F., and M.v.Z. analyzed data and wrote the manuscript.

## Data and Materials Availability

Data and materials are available upon request.

## Supplementary Figures

**Supplementary Figure 1.**
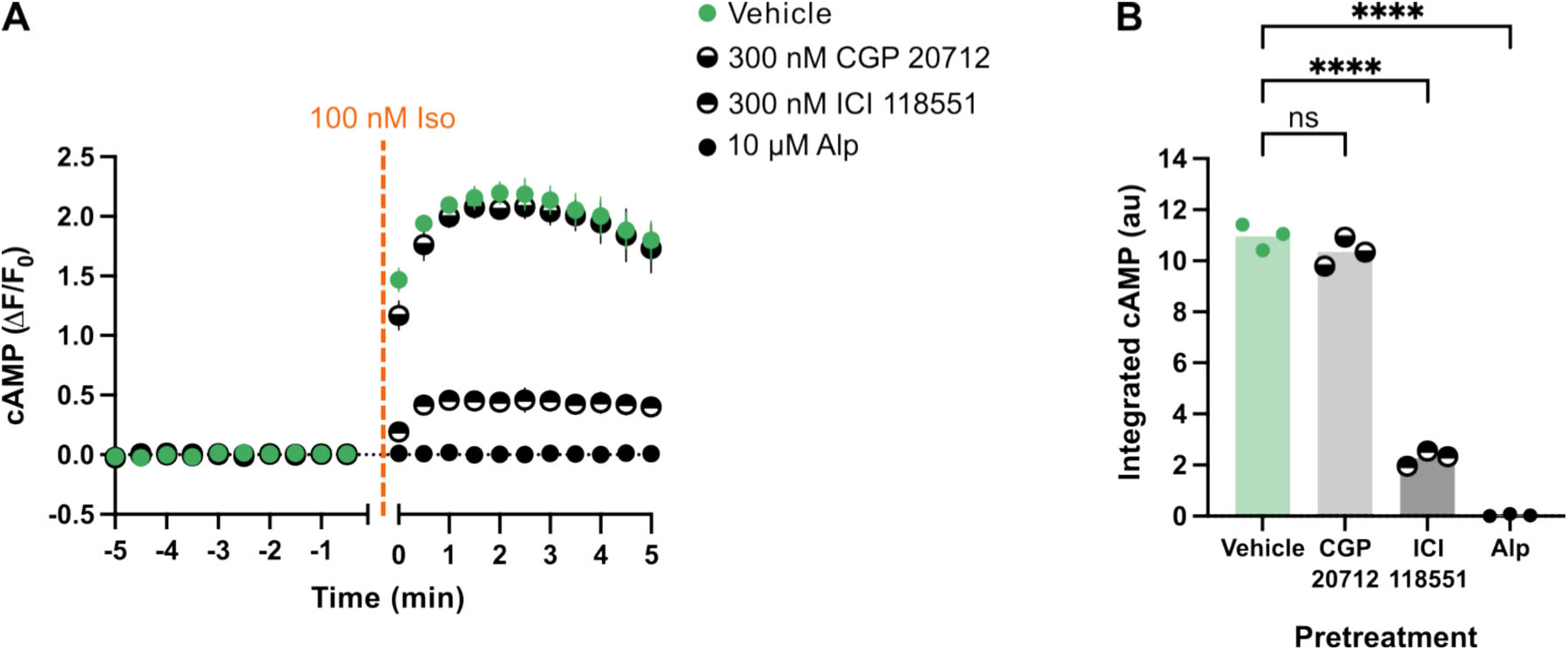
β2AR is the predominant GPCR mediating Iso-stimulated cAMP activation in HEK293. (A) Iso-induced cAMP responses in HEK293 cells pretreated for 30 minutes with vehicle or βAR antagonists: 300 nM CGP 20712 (β1AR selective), 300 nM ICI 118551 (β2AR selective), or 10 μM alprenolol (Alp, non-selective). cAMP was measured by a fluorescent cAMP biosensor, and cells were stimulated with 100 nM Iso after a 5 minute baseline reading. N = 3 independent experiments, and error bars = S.D. (B) Integrated Iso-induced cAMP from (A), calculated as an area under the curve for timepoints after Iso addition. Significance was determined using an ordinary one-way ANOVA with Dunnett’s multiple comparisons test.

**Supplementary Figure 2.**
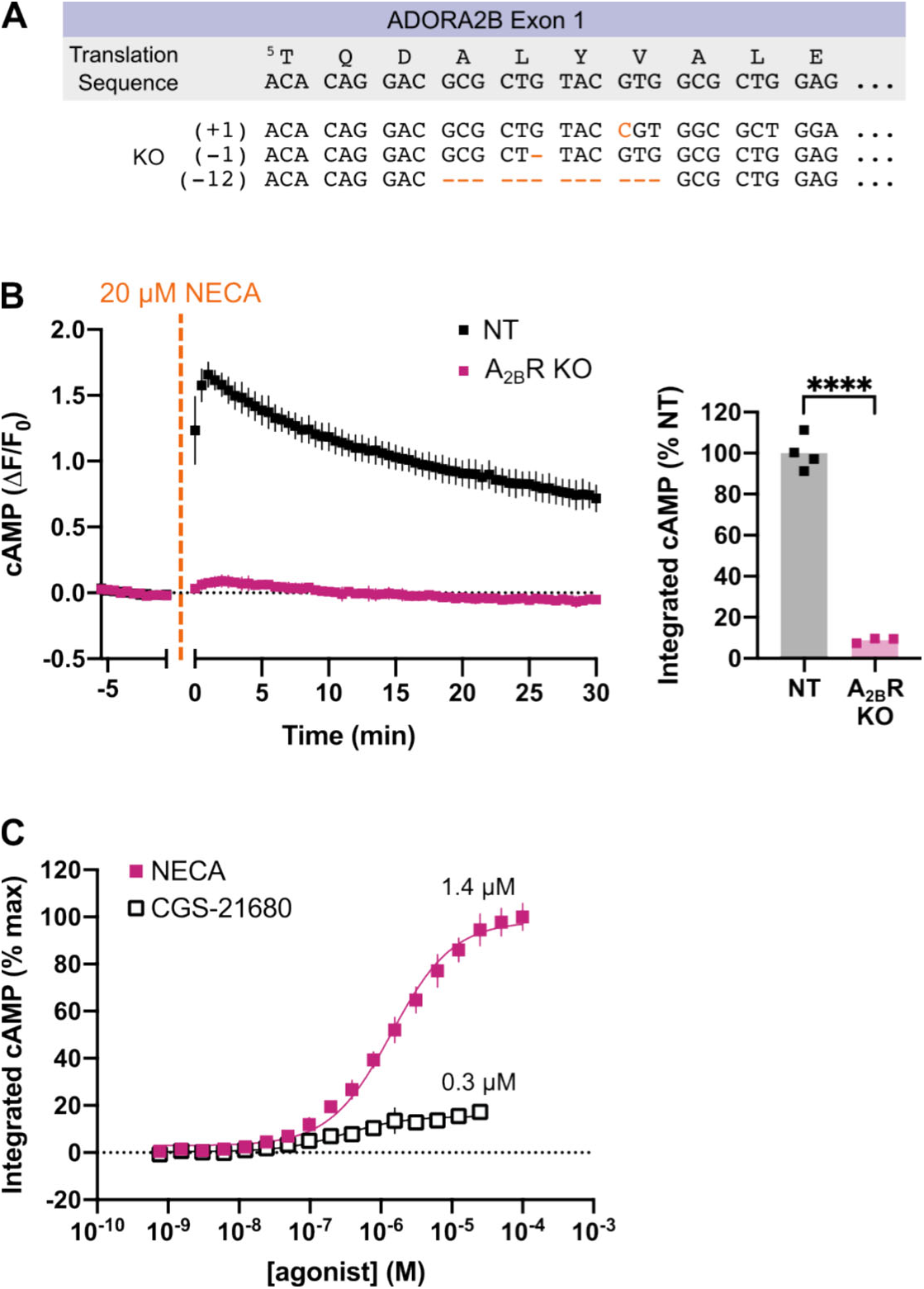
A_2B_R is the primary Gs-coupled adenosine receptor subtype endogenously expressed in HEK293. (A) *ADORA2B* genetic modifications in the A_2B_R KO cell line. (B) NECA-induced cAMP responses in control (NT) or A_2B_R KO cells, as measured by a fluorescent cAMP biosensor. Integrated cAMP was calculated as an area under the curve for timepoints after NECA addition. N = 4 (NT) or N = 3 (A_2B_R KO) independent experiments. Significance was determined using an unpaired t-test. (C) Integrated cAMP responses in wild type HEK293 cells upon stimulation with NECA or the A_2A_R-selective agonist CGS-21680. Data were normalized to the maximum NECA response and fit to a three-parameter dose-response curve. N = 3 independent experiments, and error bars = S.D.

**Supplementary Figure 3.**
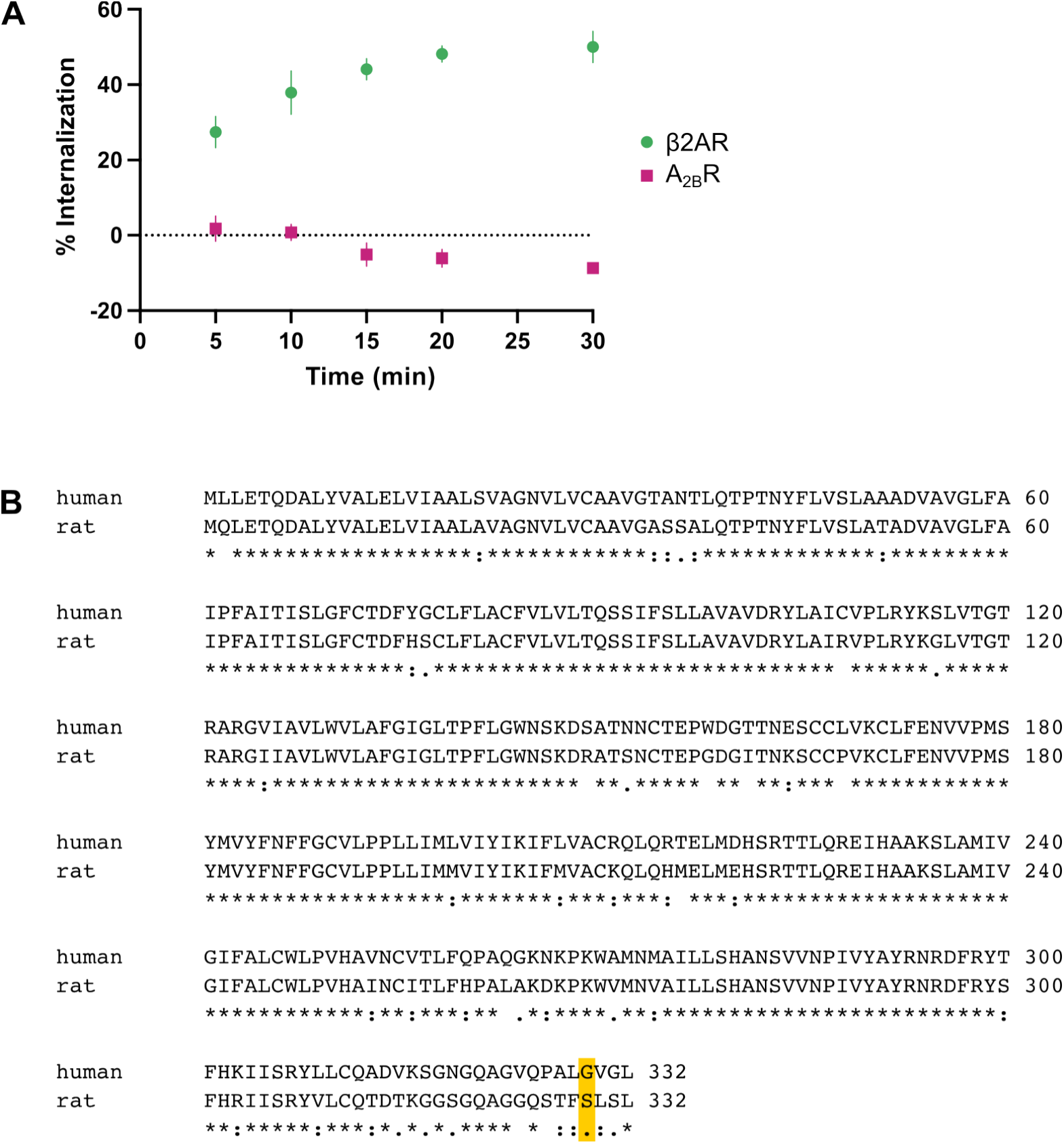
FLAG-A_2B_R does not undergo agonist-induced internalization. (A) Time course of agonist-induced internalization of FLAG-ꞵ2AR and FLAG-A_2B_R, as measured by flow cytometry. Receptors were stimulated with agonist (100 nM Iso or 20 μM NECA) for the noted times, and receptors remaining on the surface were labeled with M1 anti-FLAG antibody labeled with Alexa Fluor 647. N = 3 independent experiments, and error bars = S.D. (B) Sequence alignment of the human (Uniprot P29275) and rat (Uniprot P29276) A_2B_R, with rat S329 highlighted. Alignment made using Clustal Omega (*45*).

**Supplementary Figure 4.**
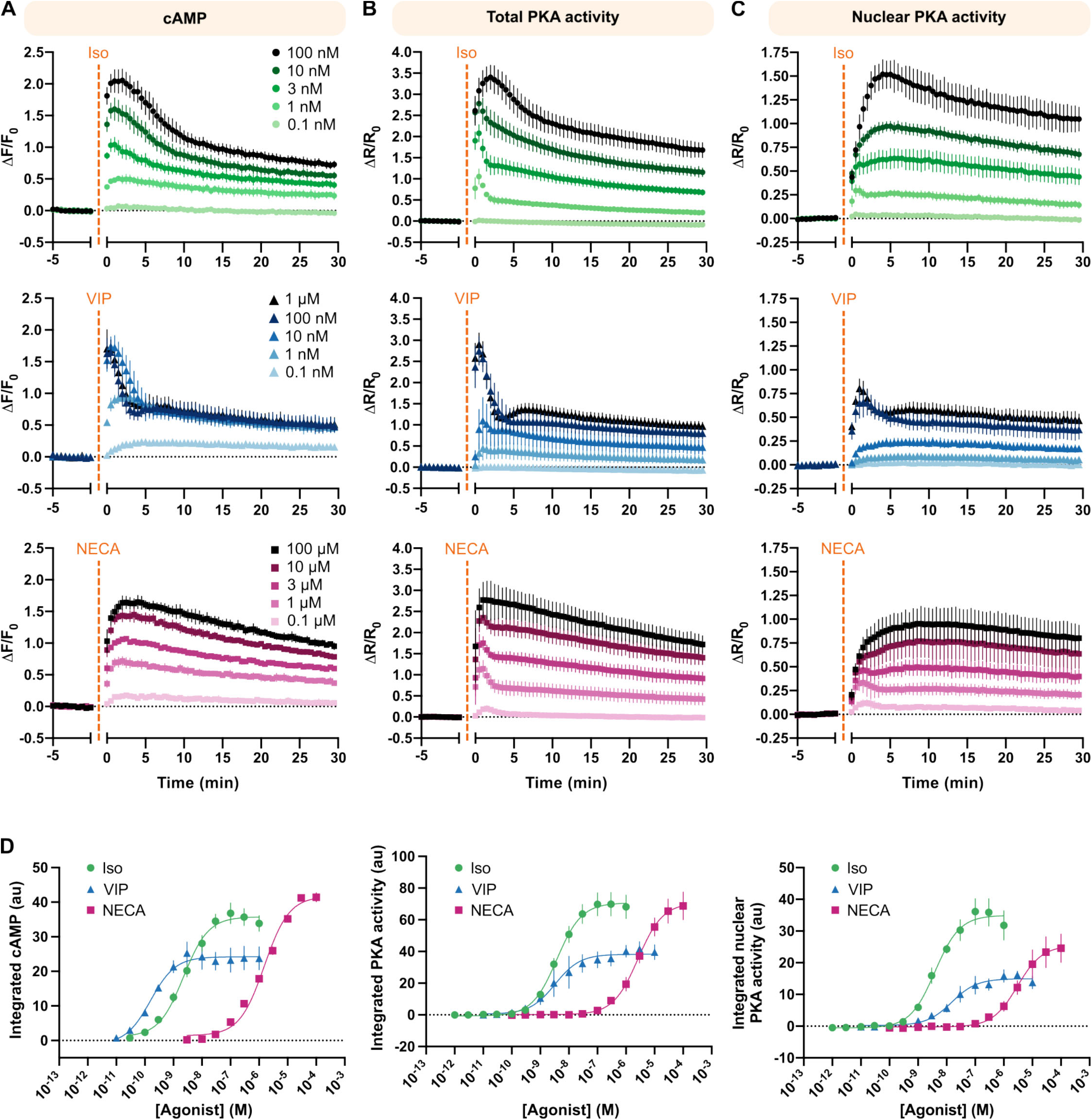
Biosensor dose-response curves in HEK293 cells. (A-C) Fluorescence time courses of biosensors for total intracellular cAMP (A), total cellular PKA activity (B), and nuclear PKA activity (C) upon stimulation of HEK293 cells with Iso (green circles), VIP (blue triangles), or NECA (pink squares) at the concentrations indicated. (D) Integrated signaling, calculated as the area under the curve after agonist addition, across doses of Iso, NECA, and VIP for all three biosensors. Corresponding time courses for a subset of data are shown in (A). Three-parameter dose-response curve fits are shown, and EC_50_s are listed in Supplementary Table 1. For all panels, N = 2 (VIP cAMP at 1 μM and 10 pM) or N = 3 (all others) independent experiments, and error bars = S.D.

**Supplementary Figure 5.**
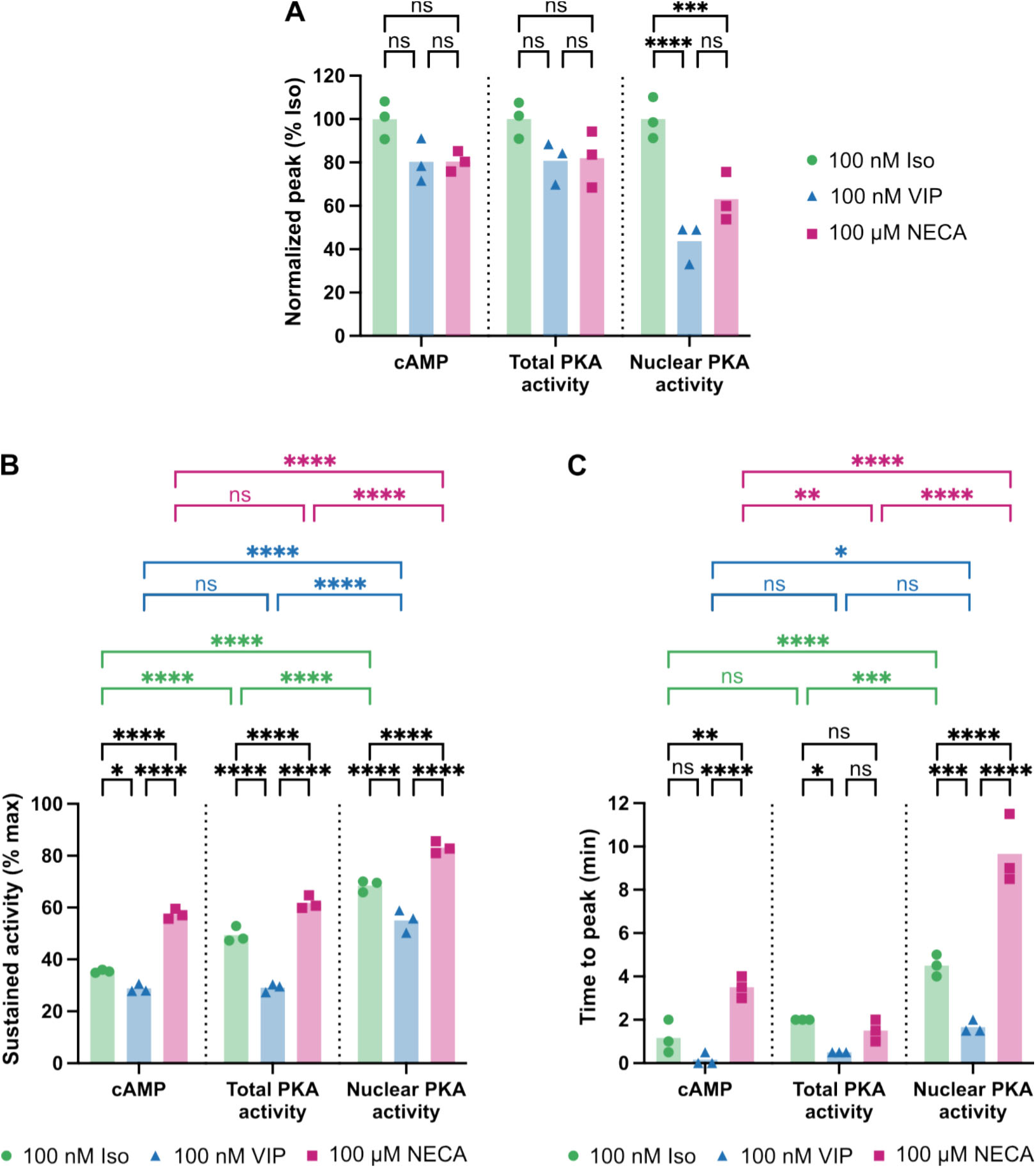
Quantification of cAMP, total PKA activity, and nuclear PKA activity responses to saturating doses of agonist. (A) Peak responses upon stimulation with 100 nM Iso (green circles), 100 nM VIP (blue triangles) or 100 μM NECA (pink squares), normalized to the average peak response upon Iso stimulation. (B) Sustained activity, calculated as the percent peak signal remaining 30 minutes after agonist addition, across agonists and biosensors. (C) Time to peak responses upon agonist stimulation across the three biosensors. For all panels, biosensor fluorescence time courses are in Figure 2, N = 3 independent experiments, and significance was determined using ordinary 2-way ANOVA with Tukey’s multiple comparisons test.

**Supplementary Figure 6.**
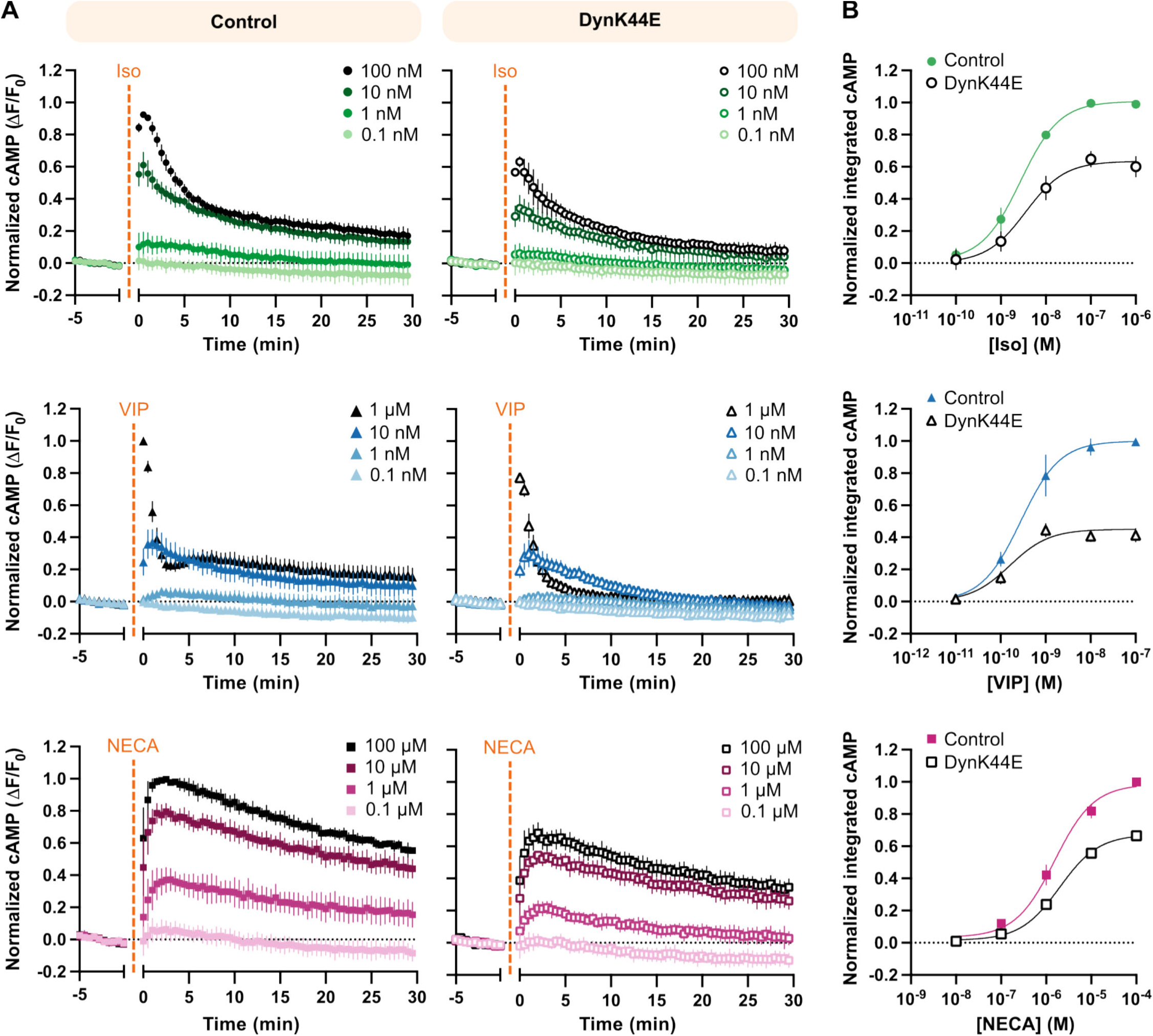
cAMP biosensor dose-response curves with and without endocytic inhibition. (A) Fluorescence time courses of a cAMP biosensor in cells expressing mCherry (control, closed shapes) or mCherry-DynK44E (DynK44E, open shapes). Cells were treated with Iso (green circles), VIP (blue triangles), or NECA (pink squares) at the indicated concentrations. Within each agonist and biological replicate, data were normalized to the max response in control cells. (B) Integrated cAMP signaling, calculated as the area under the curve after agonist addition, for cells expressing control or DynK44E. Within each agonist and biological replicate, data were normalized to the largest response in control cells. Corresponding time courses for a subset of data are shown in (A). Three-parameter dose-response curve fits are shown, and EC_50_s are listed in Supplementary Table 1. For all panels, N = 3 independent experiments, and error bars = S.D.

**Supplementary Figure 7.**
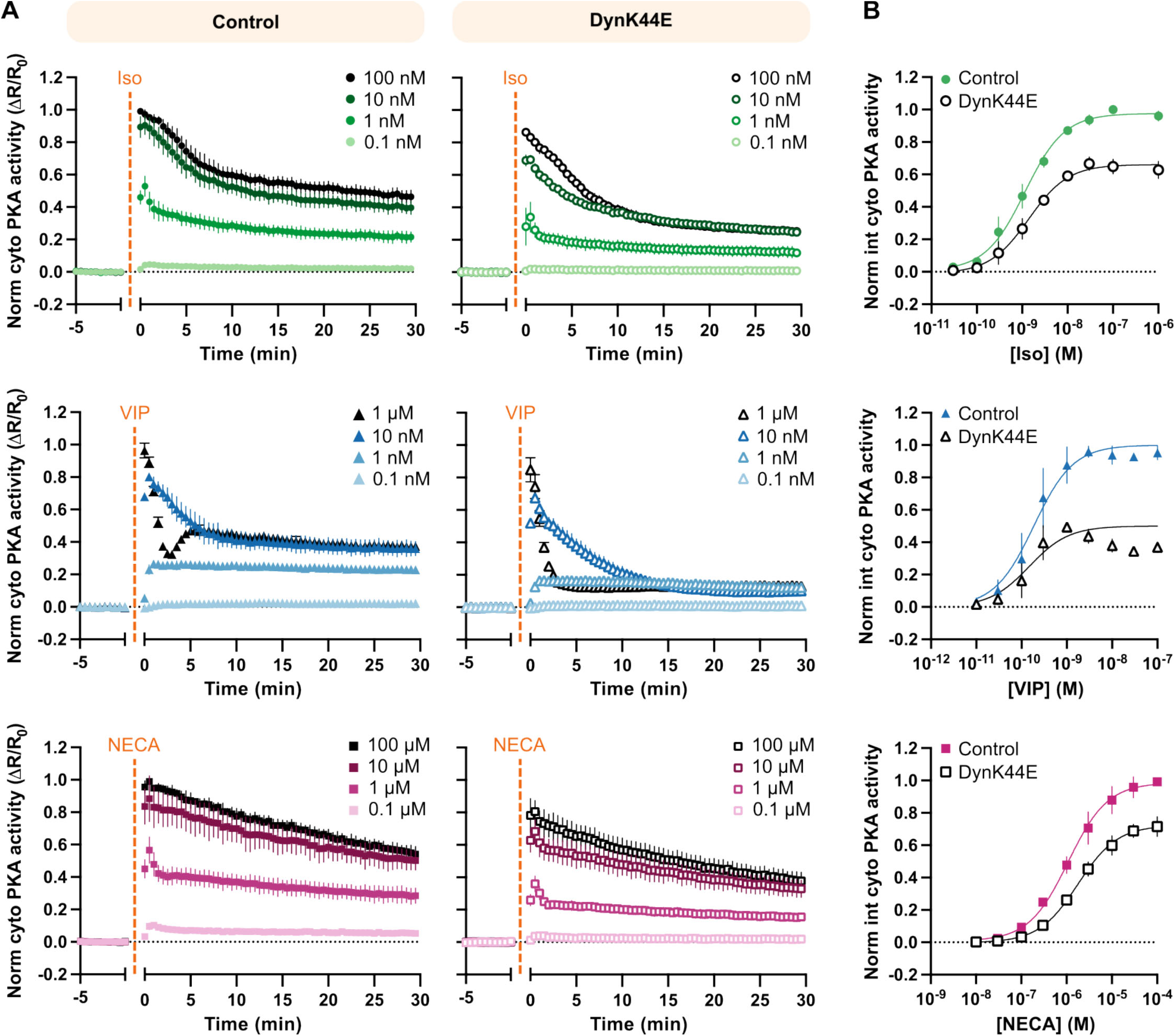
Cytoplasmic PKA activity biosensor dose-response curves with and without endocytic inhibition. (A) Fluorescence time courses of a cytoplasmic PKA activity biosensor in cells expressing mCherry (control, closed shapes) or mCherry-DynK44E (DynK44E, open shapes). Cells were treated with Iso (green circles), VIP (blue triangles), or NECA (pink squares) at the indicated concentrations. Within each agonist and biological replicate, data were normalized to the max response in control cells. (B) Integrated cytoplasmic PKA activity, calculated as the area under the curve after agonist addition, for cells expressing control or DynK44E. Within each agonist and biological replicate, data were normalized to the largest response in control cells. Corresponding time courses for a subset of data are shown in (A). Three-parameter dose-response curve fits are shown, and EC_50_s are listed in Supplementary Table 1. For all panels, N = 3 independent experiments, and error bars = S.D.

**Supplementary Figure 8.**
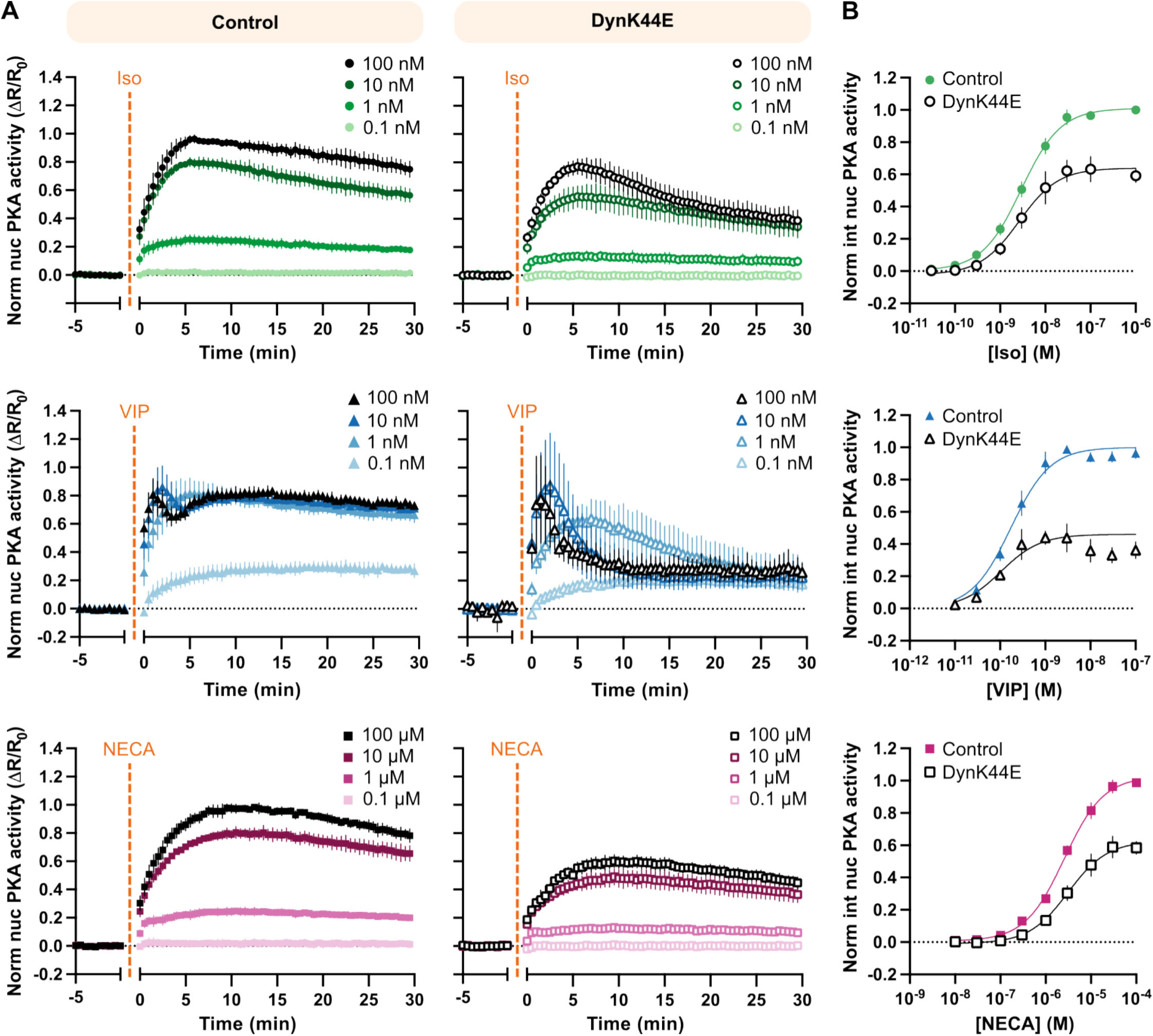
Nuclear PKA activity biosensor dose-response curves with and without endocytic inhibition. (A) Fluorescence time courses of a nuclear PKA activity biosensor in cells expressing mCherry (control, closed shapes) or mCherry-DynK44E (DynK44E, open shapes). Cells were treated with Iso (green circles), VIP (blue triangles), or NECA (pink squares) at the indicated concentrations. Within each agonist and biological replicate, data were normalized to the max response in control cells. (B) Integrated nuclear PKA activity, calculated as the area under the curve after agonist addition, for cells expressing control or DynK44E. Within each agonist and biological replicate, data were normalized to the largest response in control cells. Corresponding time courses for a subset of data are shown in (A). Three-parameter dose-response curve fits are shown, and EC_50_s are listed in Supplementary Table 1. For all panels, N = 3 independent experiments, and error bars = S.D.

**Supplementary Table 1.**
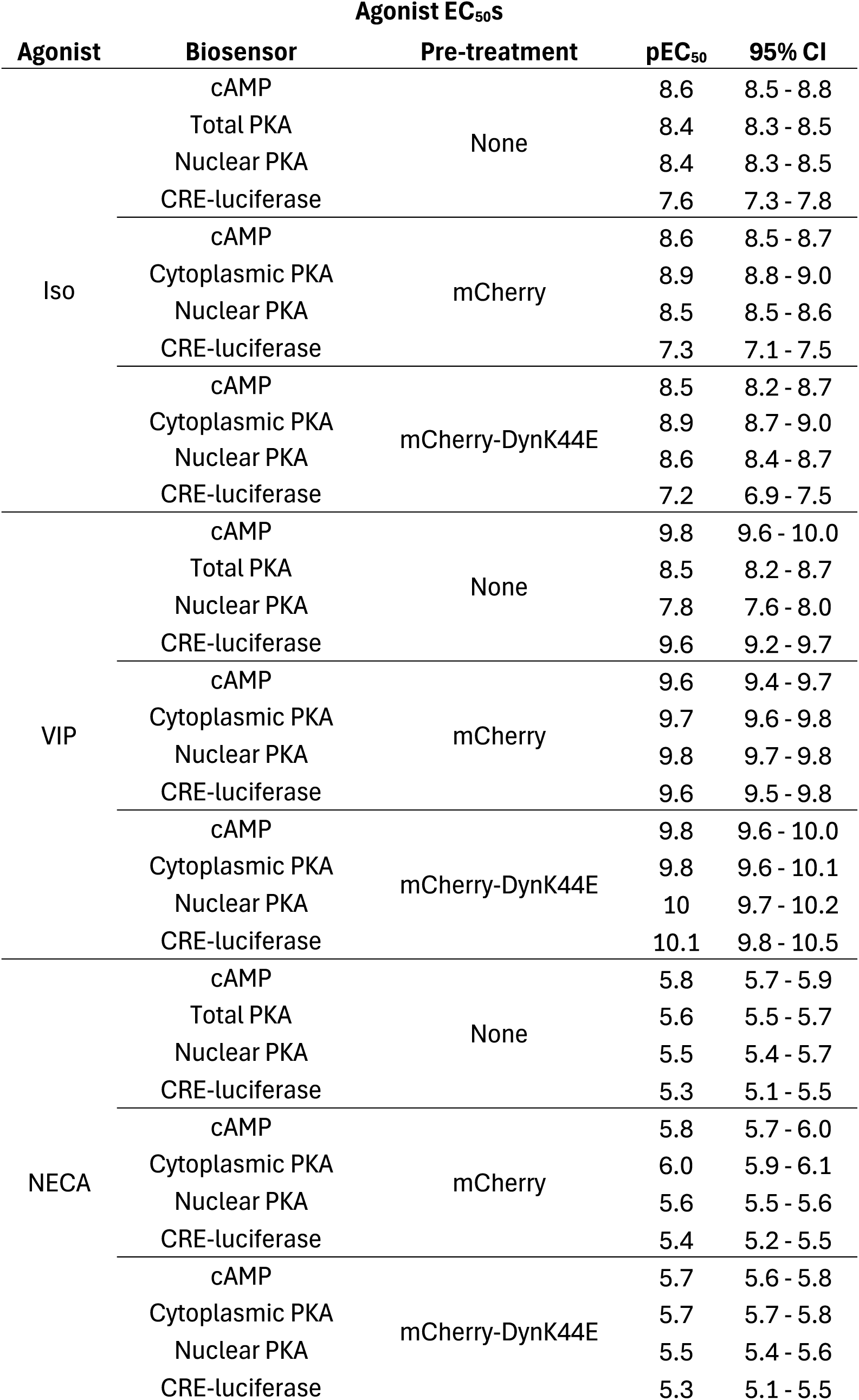
Agonist EC_50_s.

**Supplementary Table 2.**
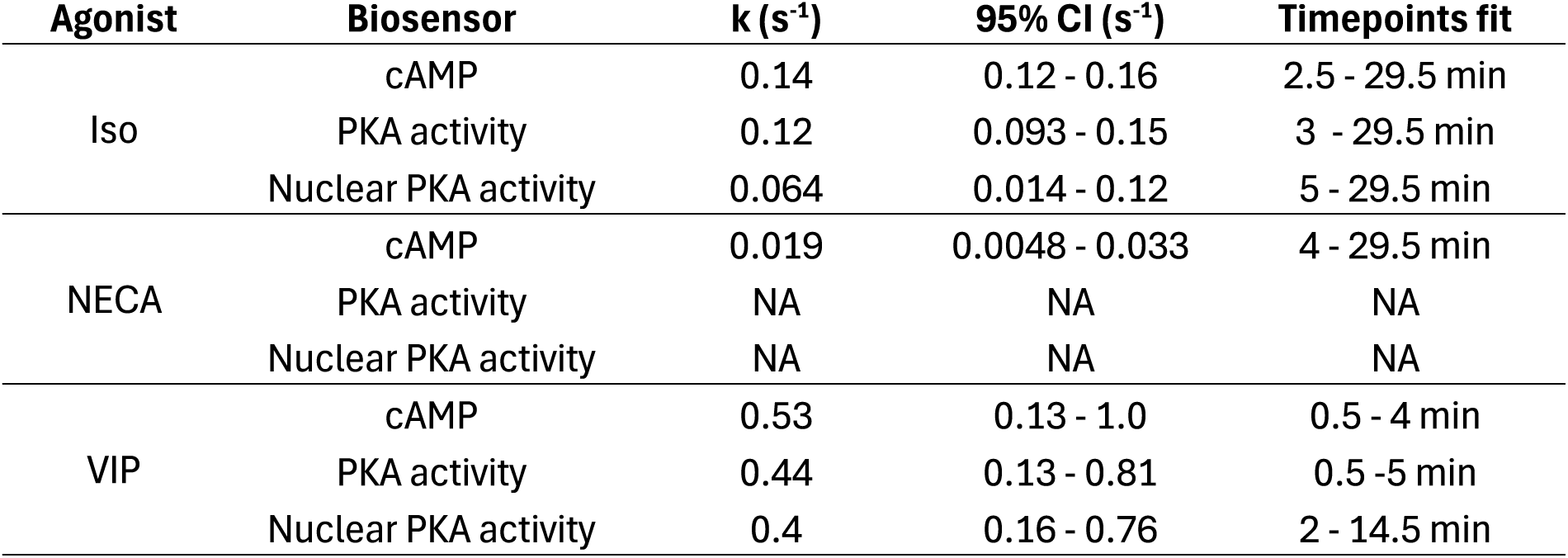
Estimated signaling decay constants. NA = could not be fit to single exponential decay.

**Supplementary Table 3.**
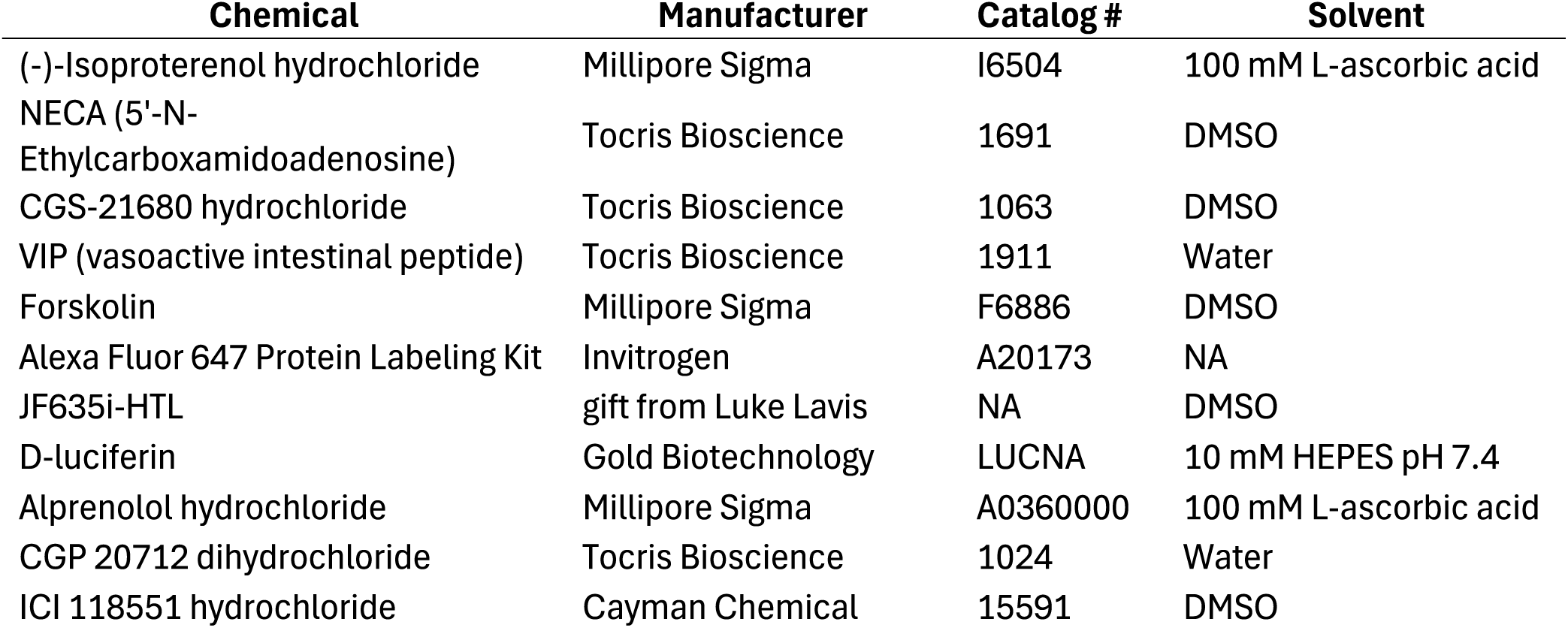
Chemicals used in this study.

| Chemical | Manufacturer | Catalog # | Solvent |
| --- | --- | --- | --- |
| (-)-Isoproterenol hydrochloride | Millipore Sigma | I6504 | 100 mM L-ascorbic acid |
| NECA (5'-N-Ethylcarboxamidoadenosine) | Tocris Bioscience | 1691 | DMSO |
| CGS-21680 hydrochloride | Tocris Bioscience | 1063 | DMSO |
| VIP (vasoactive intestinal peptide) | Tocris Bioscience | 1911 | Water |
| Forskolin | Millipore Sigma | F6886 | DMSO |
| Alexa Fluor 647 Protein Labeling Kit | Invitrogen | A20173 | NA |
| JF635i-HTL | gift from Luke Lavis | NA | DMSO |
| D-luciferin | Gold Biotechnology | LUCNA | 10 mM HEPES pH 7.4 |
| Alprenolol hydrochloride | Millipore Sigma | A0360000 | 100 mM L-ascorbic acid |
| CGP 20712 dihydrochloride | Tocris Bioscience | 1024 | Water |
| ICI 118551 hydrochloride | Cayman Chemical | 15591 | DMSO |

**Supplementary Table 4.**
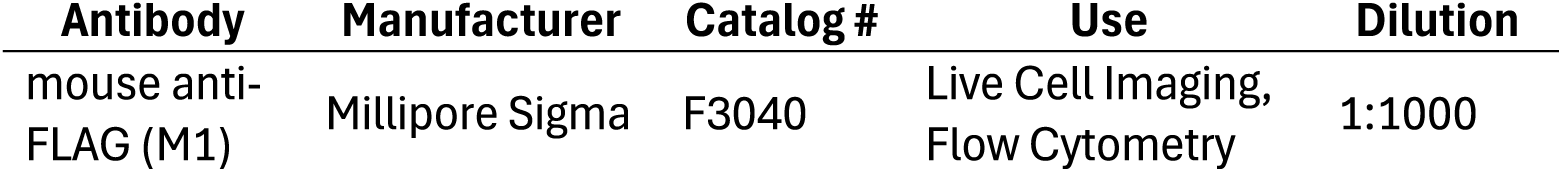
Antibodies used in this study.

**Supplementary Table 5.** DNA constructs and BacMam viruses used in this study.

| Construct | Source |
| --- | --- |
| pCMV-Dest_ExRai-AKAR2 | This study; insertion of ExRai-AKAR2 into pCMV-Dest (Invitrogen A24223) between Gateway sites. ExRai-AKAR2 was a gift from Jin Zhang (Addgene plasmid #161753) |
| pCMV-Dest-ExRai-AKAR2-NES | This study; insertion of ExRai-AKAR2-NES (30) into pCMV-Dest (Invitrogen A24223) between Gateway sites. |
| pCMV-Dest_ExRai-AKAR2-2xNLS | This study; insertion of ExRai-AKAR2-2xNLS (30) into pCMV-Dest (Invitrogen A24223) between Gateway sites. |
| pCMV-Dest-mCherry | (19) |
| pCMV-Dest-mCherry-DynK44E | (19) |
| pGL4.29[*luc2P*/CRE/Hygro] | Promega E8471 |
| pcDNA3.1-SS-FLAG-A2BR | This study; insertion of A2BR (cDNA Resource Center #ADRA2BTN00) into pcDNA3.1 at HindIII/XbaI with N-terminal sequence added: MKTIIALSYIFCLVFADYKDDDDAAAAGGGSGS |
| pcDNA3.1-SS-HaloTag-A2BR | This study; replacement of FLAG-linker (DYKDDDDAAAAGGGSGS) with HaloTag7 in pcDNA3.1-SS-FLAG-A2BR, flanked by GS (N-term) and GGGGSGS (C-term) |
| pcDNA3.1-SS-FLAG-VIPR1 | (19) |
| pcDNA-SS-FLAG-β2AR | (46) |
| pVenus-NES-Venus-miniGs | (27) |
| CMV-cADDIS Green Up | Montana Molecular #U0200G |

